# APC/C prevents non-canonical order of cyclin/CDK activity to maintain CDK4/6 inhibitor-induced arrest

**DOI:** 10.1101/2023.11.09.566394

**Authors:** Brandon L Mouery, Eliyambuya M Baker, Christine A Mills, Laura E Herring, Dalia Fleifel, Jeanette Gowen Cook

**Affiliations:** Curriculum in Genetics and Molecular Biology. The University of North Carolina at Chapel Hill. Chapel Hill, NC 27599, USA; Department of Biochemistry and Biophysics, The University of North Carolina at Chapel Hill. Chapel Hill, NC 27599; UNC Proteomics Core Facility, Department of Pharmacology. The University of North Carolina at Chapel Hill. Chapel Hill, NC 27599, USA; Lineberger Comprehensive Cancer Center. The University of North Carolina at Chapel Hill. Chapel Hill NC 27599, USA; Department of Pharmacology. The University of North Carolina at Chapel Hill. Chapel Hill NC, 27599, USA; Immuno-Oncology, Human Oncology and Pathogenesis Program, Memorial Sloan Kettering Cancer Center, New York, NY USA

**Keywords:** CDK4/6, APC/C, Cell cycle arrest, replication stress, genome instability, breast cancer, Palbociclib

## Abstract

Regulated cell cycle progression ensures homeostasis and prevents cancer. In proliferating cells, premature S phase entry is avoided by the E3 ubiquitin ligase APC/C (anaphase promoting complex/cyclosome), although the APC/C substrates whose degradation restrains G1-S progression are not fully known. The APC/C is also active in arrested cells that exited the cell cycle, but it is not clear if APC/C maintains all types of arrest. Here by expressing the APC/C inhibitor, EMI1, we show that APC/C activity is essential to prevent S phase entry in cells arrested by pharmacological CDK4/6 inhibition (Palbociclib). Thus, active protein degradation is required for arrest alongside repressed cell cycle gene expression. The mechanism of rapid and robust arrest bypass from inhibiting APC/C involves cyclin-dependent kinases acting in an atypical order to inactivate RB-mediated E2F repression. Inactivating APC/C first causes mitotic cyclin B accumulation which then promotes cyclin A expression. We propose that cyclin A is the key substrate for maintaining arrest because APC/C-resistant cyclin A, but not cyclin B, is sufficient to induce S phase entry. Cells bypassing arrest from CDK4/6 inhibition initiate DNA replication with severely reduced origin licensing. The simultaneous accumulation of S phase licensing inhibitors, such as cyclin A and geminin, with G1 licensing activators disrupts the normal order of G1-S progression. As a result, DNA synthesis and cell proliferation are profoundly impaired. Our findings predict that cancers with elevated EMI1 expression will tend to escape CDK4/6 inhibition into a premature, underlicensed S phase and suffer enhanced genome instability.

**Significance:** Appropriate stable cell cycle arrest is critical to prevent cancer. However, it is not well-understood how cells maintain arrest. It is known that arrest requires repressing proliferation-stimulating genes, but the role of targeted protein degradation is unclear. This work demonstrates that continuous degradation of cyclin A through the action of the anaphase promoting complex/cyclosome (APC/C) is required to maintain arrest induced by a cancer drug that blocks cell cycle kinase enzymes. APC/C activity is required to prevent cell cycle re-entry. Impaired APC/C activity causes arrest bypass, inefficient DNA replication, and ultimately long-term proliferation defects. These results suggest that the activity level of the APC/C in tumors may profoundly influence the response to drugs that target cell cycle kinases.

## Introduction

The balance between cell proliferation and cell cycle arrest is key to normal development and tissue homeostasis, and disruptions to this balance underlie many diseases such as cancer. One hallmark of cancer is inappropriate cell cycle entry by cells that should otherwise remain arrested (1). Several targeted cancer therapies reinforce cell cycle arrest by inhibiting specific pathways that drive progression from quiescence through G1 phase and into S phase (2). The success of these approaches relies on knowledge of the cellular systems that maintain arrest. However, despite considerable understanding of the molecular mechanisms that drive active cell cycle progression, how cells maintain arrest is less well-understood. Arrested cells have low activity in the pathways that stimulate DNA replication and mitosis, but it is clear that inhibitory mechanisms also actively restrain inappropriate cell cycle entry (3–5). Thus, arrest is characterized by both the absence of proliferation-promoting activities *and* the induction of proliferation-inhibiting activities.

One mechanism of cell cycle arrest is active repression of genes whose products promote cell cycle entry. The retinoblastoma tumor suppressor protein, RB, binds to the activator E2F family of transcription factors (E2F1, E2F2, E2F3a), thus preventing their transactivation activity and maintaining low expression of genes required for proliferation (6, 7). Upon pro-proliferative signaling, RB is phosphorylated and inactivated by cyclin-dependent kinase (CDK) activity. RB inactivation leads to its dissociation from E2F, which allows E2F to upregulate the genes required for S phase entry. In proliferating cells, cyclin-dependent kinases 4 and 6 (CDK4/6) in complex with cyclin D and CDK2 in complex with cyclin E are responsible for RB hyperphosphorylation during G1 phase (8, 9). Increased E2F activity is critical for cell proliferation, underscored by the frequency of RB/E2F dysregulation in cancer (10, 11) and the effectiveness of CDK4/6 inhibitors at blocking proliferation through RB (12).

In parallel with RB-mediated transcriptional repression of proliferation-stimulating genes, many cell cycle proteins are targeted for active ubiquitin-mediated degradation by the Anaphase Promoting Complex or Cyclosome (APC/C) in G1 phase and in arrested cells. The APC/C is a large E3 ubiquitin ligase most well-known for its role in mitotic cyclin degradation to promote mitosis completion (13). However, once cells exit mitosis to G1 phase, the APC/C remains active and switches substrate adapters from CDC20 in mitosis to CDH1 in G1 to target pro-proliferative proteins for degradation (14). Although CDH1 is dispensable for proliferation, its loss in both human cells and mouse embryonic fibroblasts causes premature S phase entry, prolonged S phase, and the accumulation of replication stress and genomic/chromosomal aberrations (15–23). Additionally, mice heterozygous for CDH1 are more susceptible to spontaneous tumor formation, indicating that CDH1 is a haploinsufficient tumor suppressor protein (18). These findings highlight the critical role of APC/C^CDH1^ activity in restraining inappropriate S phase entry and preserving genomic integrity.

Despite the knowledge that APC/C^CDH1^ activity contributes to G1/S timing control, the relevant substrate(s) whose degradation delays the onset of S phase are not fully known. One study found that cells expressing an APC/C resistant mutant of cyclin A initiate DNA replication early, supporting the notion that cyclin A must be degraded during G1 to prevent untimely S phase entry (19). Conflicting with these data, however, another study of the short G1 in CDH1-depleted cells found that cyclin A abundance was not important for setting the length of G1 (whereas cyclin E abundance was), suggesting that cyclin A is not rate-limiting for S phase entry (20). Although cyclin E is not a direct APC/C substrate, elevated levels of Skp2 in CDH1-depleted cells led to increased p27 degradation, and thus reduced the amount of cyclin E necessary to stimulate CDK2-mediated S phase entry. Given these conflicting findings, the exact mechanism(s) by which APC/C^CDH1^ restrains unscheduled S phase entry remains unknown.

Moreover, it remains unclear if APC/C^CDH1^ activity is similarly required to maintain cell cycle arrests such as quiescence (G0), senescence, or arrests induced by pharmacological inhibitors. Some studies have shown that CDH1 depletion *<u>prior</u>* to an arrest-promoting signal (serum withdrawal, CDK4/6 inhibition) prevents arrest (19, 24). However, little is known about whether APC/C^CDH1^ activity remains essential *after* the arrested state has been established. To address this question, we investigated the effects of inhibiting APC/C activity in cells arrested by the CDK4/6 inhibitor, Palbociclib (“CDK4/6i”). We demonstrate that APC/C activity is essential to maintain G1 arrest induced by CDK4/6 inhibition and that APC/C inhibition in arrested cells results in escape from arrest mediated by non-canonical order of cyclin/CDK activity. We provide evidence that cyclin A degradation is absolutely required for arrest and that cyclin B degradation inhibits cyclin A accumulation. APC/C-mediated degradation of both cyclin A and cyclin B prevents RB phosphorylation and E2F-dependent transcription. Furthermore, we show that cells bypassing arrest enter S phase with reduced origin licensing and display reduced DNA synthesis rates and long-term proliferation defects. Collectively, our results highlight the essential role of APC/C activity in maintaining cell cycle arrest and preserving proliferative fitness, and we suggest that the status of APC/C activity in cancers may profoundly influence the response to clinical CDK4/6 inhibition.

## Results

### APC/C activity is essential to maintain arrest induced by CDK4/6 inhibition

To address whether APC/C activity is necessary to restrain DNA replication and cell cycle progression during an established cell cycle arrest, we first arrested cells with the CDK4/6 inhibitor, Palbociclib, and then inhibited the APC/C (Fig. 1*A*). To inhibit APC/C activity, we used a previously-described non-transformed retinal pigmented epithelial (RPE1) cell line in which the natural APC/C inhibitor protein, EMI1, is induced by doxycycline (“dox”) (25). We analyzed the cell cycle distribution of cells treated with Palbociclib for 48 hours with or without APC/C inhibition during the final 24 hours by flow cytometry. We included the nucleotide analog, EdU, in the final hour as a measure of active DNA synthesis (see Methods). This analysis confirmed that cells were completely arrested with G1 DNA content (blue) after 24 hours of Palbociclib treatment and further, that control cells remained arrested after an additional 24 hours, as determined by DNA content and no EdU incorporation (Fig. 1*B* and Fig. S1). In contrast, APC/C inhibition during the final 24 hours led to a marked increase in the percentage of cells in either S phase (EdU-positive, orange) or G2/M phase (4C DNA, EdU-negative, grey) and a corresponding reduction in G1 (Fig. 1*B*) (For simplicity, we refer to CDK4/6i-arrested cells as “G1,” but acknowledge that they may exhibit characteristics of quiescence (G0), senescence, and/or other distinct features.) (26). Even near-endogenous EMI1 levels induced S phase entry within 24 hours, indicating that escape from arrest does not require highly overexpressed EMI1 (Fig. S2*A-C*). EMI1 expression also modestly decreased the G1 fraction in cells that were arrested by either contact inhibition or serum starvation, revealing that APC/C activity maintains multiple forms of cell cycle arrest (Fig. S3*A-B*).

**Figure 1.**
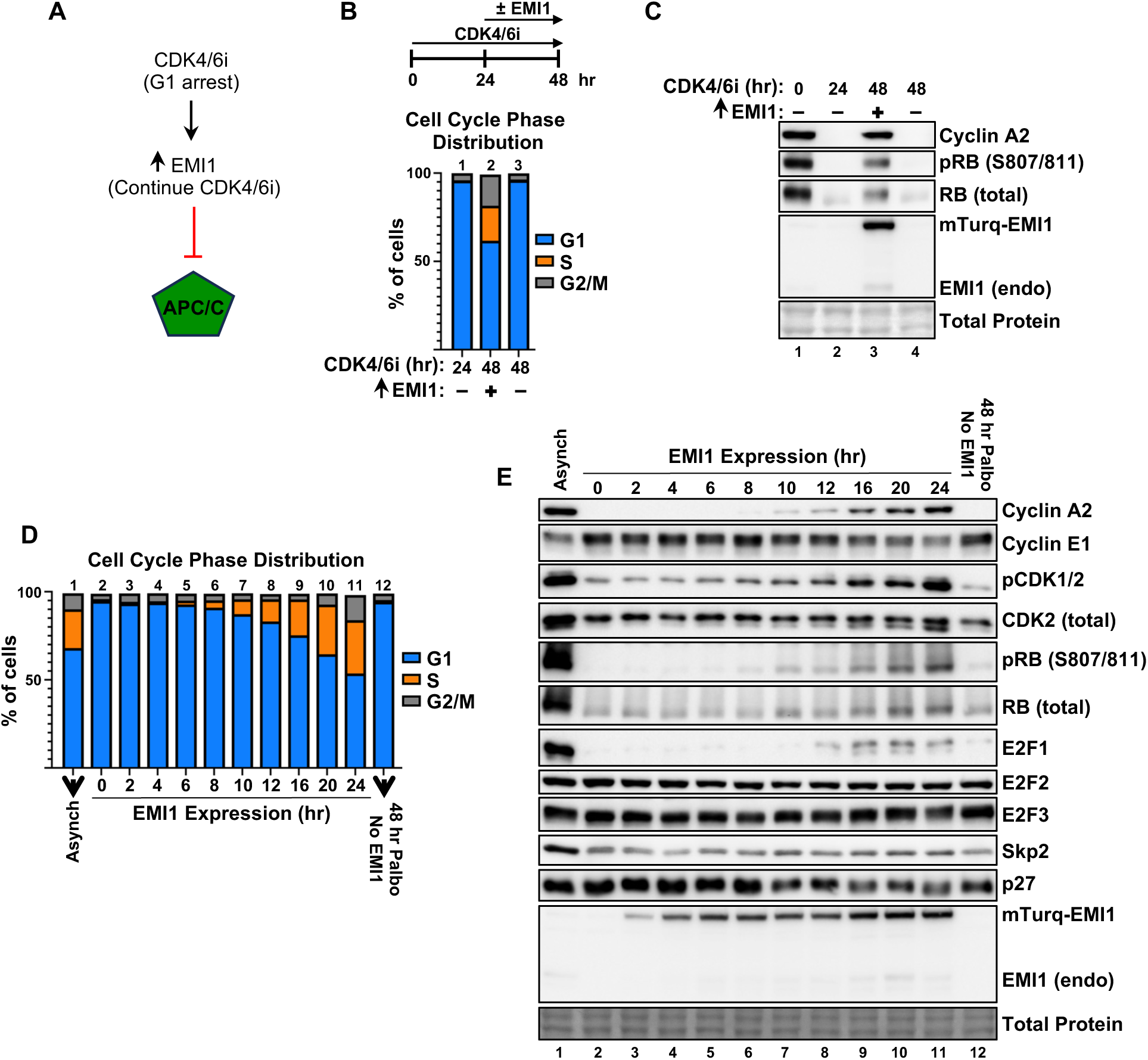
APC/C activity is essential to maintain CDK4/6 inhibitor-induced G1 arrest in RPE1 cells. **(A)** General experimental outline used in this study. Cells are arrested in Palbociclib (CDK4/6i) prior to induction of EMI1 expression to inhibit the activity of the APC/C. Cells remain in the presence of CDK4/6i throughout the experiment. **(B)** Cell cycle distribution determined by flow cytometry of RPE1 cells treated with 1 µM CDK4/6i for 48 hours ± EMI1 expression during the final 24 hours as indicated. One sample was collected after 24 hours of CDK4/6i to show that cells were fully arrested at the time of EMI1 induction. **(C)** Immunoblot of lysates from (B). An untreated proliferating sample (“Async.”) was included as a control. **(D)** Cell cycle distribution of RPE1 cells treated with 1 µM CDK4/6i for 24 hours and then induced to express EMI1 for 0-24 hours. An untreated proliferating sample (“Asynch”; column 1) was included as a control. **(E)** Immunoblot of lysates from (D). For cell cycle and immunoblots, one representative example is shown. Panels B-C are representative of more than 3 biological replicates and panels D and E are representative of 2 biological replicates.

As expected, both phosphorylated RB (pRB) and the APC/C substrate cyclin A were undetectable in arrested cells (Fig. 1*C*, compare lanes 1 and 2). Strikingly, inhibiting the APC/C in Palbociclib-arrested cells was sufficient to restore pRB and cyclin A nearly to levels in proliferating control cells (Fig 1*C*, compare lanes 1 and 3). pRB and cyclin A remained undetectable in cells that were instead treated only with Palbociclib (without EMI1 expression), confirming that their reappearance was not due to metabolism of, or rapid adaptation to, the inhibitor (Fig. 1*C*, compare lanes 3 and 4). Endogenous EMI1 (lower band in Fig. 1*C*) was undetectable in arrested cells but was restored upon inhibiting the APC/C, likely both because it is the product of an E2F-regulated gene and because at low levels EMI1 is also a substrate of the APC/C (27, 28). From these data, we conclude that continuous APC/C activity is essential to maintain G1 arrest induced by pharmacological CDK4/6 inhibition.

To better understand this phenomenon, we analyzed the effects of EMI1 expression in Palbociclib-arrested cells for times ranging from 0-24 hours. Cells began to enter S phase as early as 6-8 hours after EMI1 induction, and ~50% of cells escaped G1 arrest within 24 hours (Fig. 1*D*). To extensively characterize escaping cells, we immunoblotted for proteins involved in the G1/S transition. We found that cyclin A and pRB were first detectable around 8 hours, roughly the same time that cells began entering S phase (Fig. 1*E*, compare lane 2 to lane 6). Interestingly cyclin E levels increased rather than decreased in arrested cells (compare lane 1 to lane 2) and remained at this elevated level until declining at 16-20 hours (compare lane 2 to lanes 9 and 10), likely reflecting SCF^Fbw7^-induced degradation in S phase cells (Fig. 1*E*) (29, 30). We also found that Palbociclib treatment downregulated the activating T-loop phosphorylation of CDK2 residue T160 (compare lanes 1 and 2), which reappeared in escaping cells around 6-8 hours (Fig. 1*E*; monitored by phospho-specific antibody and by a faster-migrating band using a total CDK2 antibody). We note that this phospho-specific antibody cross-reacts with the activating T-loop phosphorylation of CDK1 residue T161, indicating that CDK1 is also unphosphorylated in arrested cells. E2F1 levels were almost undetectable in arrested cells (compare lane 1 to lane 2) and did not begin to accumulate until 12 hours after EMI1 induction (compare lane 2 to lane 8). In contrast, the other activator E2F proteins (E2F2 and E2F3) remained stable throughout the entire timecourse (Fig. 1*E*). Based on previous reports, we tested if re-accumulation of the APC/C substrate, Skp2, may drive escape from arrest. Skp2 targets p27 for degradation in late G1 phase, and premature Skp2-mediated p27 degradation and elevated CDK activity in APC/C-inhibited cells could promote S phase entry (15, 16, 31, 32). However, we observed neither an increase in Skp2 levels nor a decrease in p27 levels when we expressed EMI1 in arrested cells, suggesting that escape from arrest is not mediated by the Skp2/p27 axis (Fig. 1*E*, compare lane 2 to lanes 3-12).

In addition to its role in inhibiting the APC/C, EMI1 also contains an F-box domain through which it assembles into a SKP1-CUL1-F-box protein (SCF) ubiquitin ligase complex and recruits substrates for degradation (33, 34). We thus tested if escape from arrest is through a non-canonical EMI1 F-box-dependent mechanism by expressing an EMI1 variant lacking the F-box domain (34); this variant induced escape from G1 arrest similar to wild-type EMI1 (Fig. S4*A-B*). We conclude that escape from CDK4/6i-induced arrest following EMI1 expression is due to loss of APC/C activity.

### RB maintains CDK4/6 inhibition-induced arrest

It is well-established that loss of RB promotes resistance to CDK4/6 inhibitor treatment (12). However, it has been reported that CDK4/6 inhibition paradoxically leads to RB degradation (35, 36), and we confirmed that the total levels of RB were indeed strongly downregulated by Palbociclib treatment, although cells remained arrested (Fig. 1*C*, compare lanes 1 and 2). It was thus unclear whether RB is necessary to *maintain* a CDK4/6i-induced arrest following its establishment. To address this question, we arrested cells in Palbociclib, then treated with siRNA to deplete the residual RB protein. RB depletion was sufficient to cause re-accumulation of cyclin A and promote S phase entry (Fig. 2*A*), revealing that, despite its very low levels, RB is required to maintain arrest even after arrest is established (Fig. 2*B*, compare lanes 2 and 4). The re-appearance of pRB in APC/C-inhibited cells (Fig. 1*C*) prompted us to also test for the re-expression of several representative E2F-dependent genes. Palbociclib treatment strongly repressed expression of each gene to 20 percent or less of expression in proliferating control cells (Fig. 2*C*, brown). APC/C inhibition induced re-expression of each analyzed gene 2-20-fold relative to Palbociclib treatment alone (Fig. *2C*, green).

**Figure 2.**
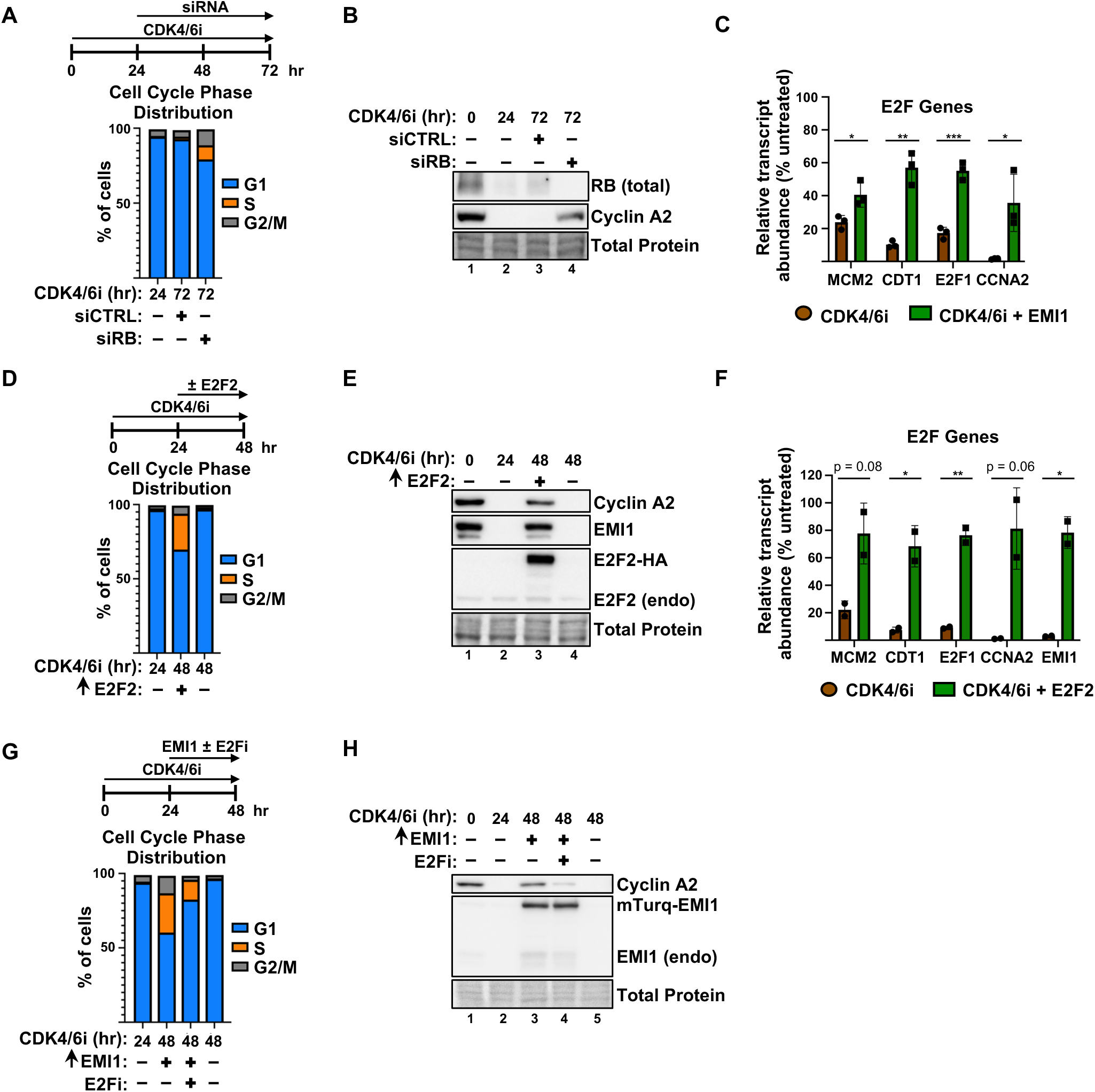
APC/C activity enforces G1 arrest by maintaining RB-dependent E2F gene repression. **(A)** Cell cycle distribution of RPE1 cells treated with 1 µM CDK4/6i for 72 hours ± control siRNA (“CTRL”) or siRNA targeting RB (“siRB”) during the final 48 hours as indicated. One sample was collected after 24 hours of CDK4/6i to show that cells were fully arrested at the time of siRNA transfection. **(B)** Immunoblot of lysates from (A). An untreated proliferating sample (lane 1) was included as a control. **(C)** Relative mRNA abundance of E2F-regulated transcripts from RPE1 cells treated with 1 µM CDK4/6i for 48 hours ± EMI1 expression during the final 24 hours as indicated. Transcript abundances are normalized to a proliferating control sample. Error bars display the mean ± SD of n = 3 biological replicates. Significance determined by unpaired t test (*, P ≤ 0.05; **, P ≤ 0.01; ***, P ≤ 0.001). **(D)** Cell cycle distribution of RPE1 cells treated with 1 µM CDK4/6i for 48 hours ± E2F2 overproduction during the final 24 hours as indicated. One sample was collected after 24 hours of CDK4/6i to show that cells were fully arrested at the time of E2F2 induction. **(E)** Immunoblot of lysates from (D). **(F)** Relative mRNA abundance of E2F-regulated transcripts from cells treated as in (D). Transcript abundances are normalized to a proliferating control sample. Error bars display the mean ± SD of n = 2 biological replicates. Significance determined by unpaired t test (*, P ≤ 0.05; **, P ≤ 0.01). **(G)** Cell cycle distribution of RPE1 cells treated with 1 µM CDK4/6i for 48 hours ± EMI1 expression in the presence or absence of 20 µM E2F inhibitor (HLM006474) during the final 24 hours as indicated. One sample was collected after 24 hours of CDK4/6i to show that cells were fully arrested at the time of EMI1 induction. **(H)** Immunoblot of lysates from (G). For cell cycle and immunoblots, one representative example is shown. Panels A-B are representative of 3 biological replicates and panels D-E and G-H are representative of 2 biological replicates.

To determine if low E2F activity is required for CDK4/6i-induced arrest, as opposed to other functions of RB (37, 38), we directly manipulated E2F in arrested cells. We overproduced the activator E2F, E2F2, which we chose to avoid the apoptosis-inducing activity of E2F1 (39). E2F2 overexpression did indeed induce escape from arrest (Fig. 2*D*). It also induced cyclin A expression and bypassed repression of representative Palbociclib-repressed genes (Fig 2*E-F*). These data demonstrate that increased E2F activity can bypass CDK4/6i-induced arrest in addition to the previously-reported ability to bypass quiescence (40).

Having established that E2F activity is *sufficient* to promote S phase entry in Palbociclib-arrested cells (Fig. 2*D*), we next tested if E2F activity is *necessary* for escape in APC/C-inhibited cells. In our timecourse analysis, E2F1 was the only activator E2F protein that was downregulated in arrested cells (Fig. 1*E*). We thus surmised that loss of E2F1 is important for arrest and that its re-accumulation upon APC/C inhibition promotes E2F-dependent gene expression and S phase entry. Blocking E2F1 re-accumulation with siRNA reduced the percentage of cells escaping arrest relative to controls (Fig S5*A-B*). However, despite the modest reduction, we still observed considerable S phase entry in E2F1-depleted cells. Given that the other activator E2Fs remained abundant in arrested cells (Fig. 1*E*), we next hypothesized that an increase in their activity contributes to escape. Consistent with this hypothesis, we found that inhibiting all E2F activity using a pan-E2F small molecule inhibitor at low doses (41) also reduced the rate of S phase entry (Fig. 2*G-H*). Altogether, we conclude that APC/C activity is required to maintain RB-dependent repression of E2F-regulated genes, and that this repression is necessary for sustained arrest by CDK4/6 inhibition.

### Premature cyclin B accumulation in cells bypassing arrest

We were particularly interested in understanding the molecular mechanism of escape from arrest when we inhibited APC/C activity. However, our timecourse analysis of candidate proteins (Fig. 1*E*) did not identify any that significantly accumulated before cells had already started to enter S phase. This circumstance made distinguishing which changes were a *cause* rather than a consequence of S phase entry challenging. Furthermore, our analysis was limited to pre-selected proteins known to regulate S phase entry. To identify all potential APC/C substrates driving escape from arrest, we performed proteomic profiling of cells escaping arrest at both 8 and 20 hours after inducing EMI1. We chose the 20-hour timepoint because at this time a significant number of cells have already escaped arrest (Fig. 1*D*) and we anticipated that - while we may identify many cell cycle-regulated non-substrates and S phase proteins - most relevant APC/C substrates would be detectable. In contrast, we chose the 8-hour timepoint because it is near the time when cells just begin to escape arrest (Fig. 1*D*). We expected to detect fewer changes overall but reasoned that those changes would include the APC/C substrates *causing* escape. We prioritized proteins that were first significantly downregulated after 24 hours of Palbociclib treatment (relative to asynchronous/proliferating control cells) and then significantly upregulated following APC/C inhibition (relative to 24 hours Palbociclib treatment alone) as those most likely to be *bona fide* APC/C substrates directly involved in arrest bypass. Of the proteins upregulated at the 8-hour timepoint, 15/22 (68%) met these criteria. Similarly, 65/125 (52%) of the proteins upregulated at the 20-hour timepoint also met these criteria (Fig. 3*A* and Supp Table 1; proteins meeting analysis criteria colored in red in Fig 3*A*).

**Figure 3.**
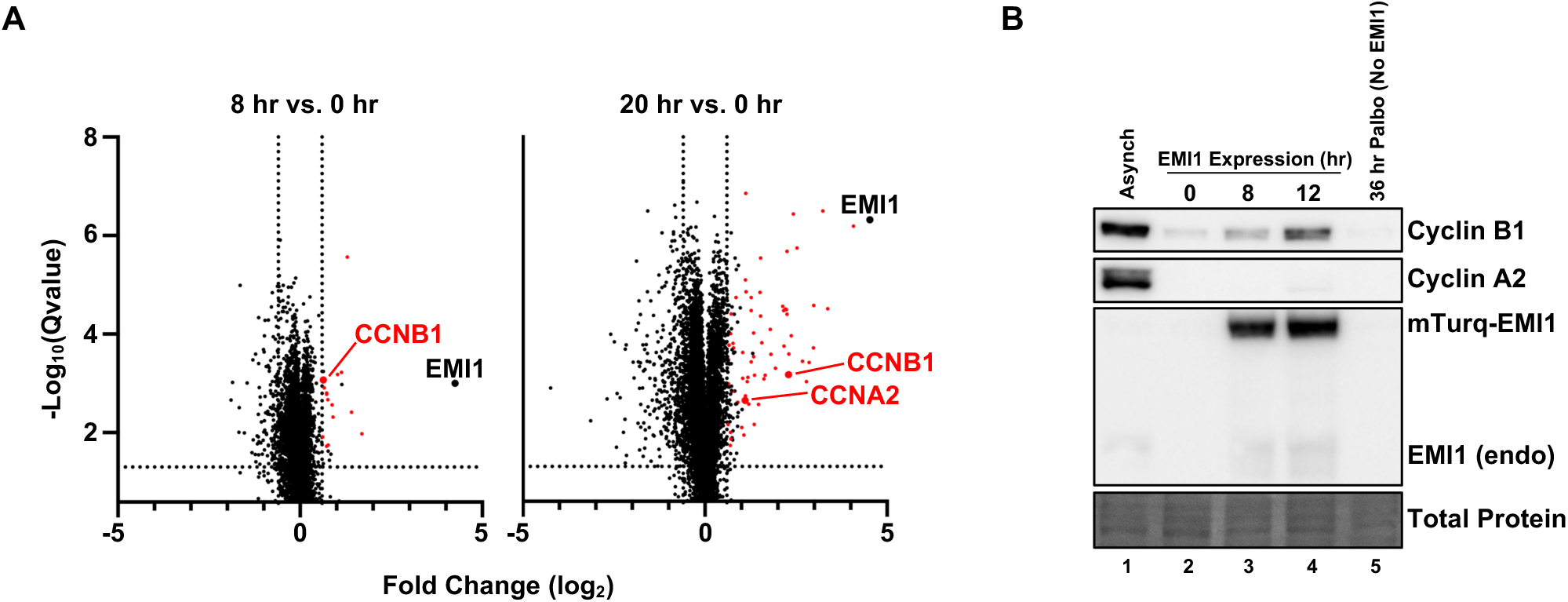
Premature cyclin B accumulation in cells bypassing CDK4/6i-induced arrest. **(A)** Volcano plot showing changes in protein abundance following APC/C inhibition in CDK4/6-inhibited cells. RPE1 cells were first arrested in 1 µM CDK4/6i for 24 hours (t = 0 hour) and then induced to express EMI1 for either 8 hours or 20 hours and analyzed by LC-MS/MS. The left plot shows changes in protein abundance after 8 hours of EMI1 expression (relative to the t = 0 hour timepoint) and the right plot shows changes in protein abundance after 20 hours of EMI1 expression (relative to the t = 0 hour timepoint). Colored in red are proteins that were significantly downregulated after 24 hours of CDK4/6i (relative to a proliferating control sample) and then significantly upregulated following EMI1 expression (relative to the t = 0 hour timepoint). All sample conditions performed in technical triplicate. **(B)** Immunoblot of lysates from RPE1 cells treated with 1 µM CDK4/6i for 24 hours and then induced to express EMI1 for 0-12 hours as indicated. One representative example of 2 biological replicates is shown.

Most strikingly, cyclin B accumulated significantly within the first 8 hours of APC/C inhibition, whereas cyclin A was only significantly upregulated in the 20-hour sample (Fig. 3*A*). We validated this result by immunoblot, observing cyclin B increasing after 8 hours of EMI1 expression, whereas cyclin A was still undetectable at this time (Fig. 3*B*). We conclude that although both cyclin A and cyclin B are stabilized upon APC/C inhibition in arrested cells, cyclin B is stabilized with faster kinetics and is detectable prior to a substantial percentage of cells escaping arrest.

### Cyclin/CDK requirements for escape from arrest

The observation that cyclin B accumulates faster than cyclin A and very close to the time of S phase entry in cells escaping from arrest raised the intriguing possibility that cyclin B – which is normally associated with and most active during G2 and M phases - is important for S phase entry in this context. To test this idea, we arrested cells in Palbociclib prior to transfection with siRNAs targeting either cyclin A or cyclin B to prevent their re-accumulation during APC/C inhibition (two different siRNAs per cyclin). Surprisingly, blocking either cyclin A or cyclin B accumulation prevented S phase entry, indicating that *both* cyclins contribute to escape from arrest (Fig. 4*A-B* and S6*E-F*). Depleting either cyclin A or cyclin B in otherwise untreated proliferating cells did not induce G1 accumulation, indicating that 1) neither cyclin is required for S phase entry in normal cell cycles and 2) inability to bypass arrest is unlikely a non-specific proliferation defect caused by the siRNAs (Fig. S6*A-D*). Another APC/C substrate, geminin (42), was stabilized in both cyclin A and B-depleted cells, confirming that the APC/C was still effectively inhibited (Fig. 4*B*; compare lane 6 to 7 and lane 9 to 10).

**Figure 4.**
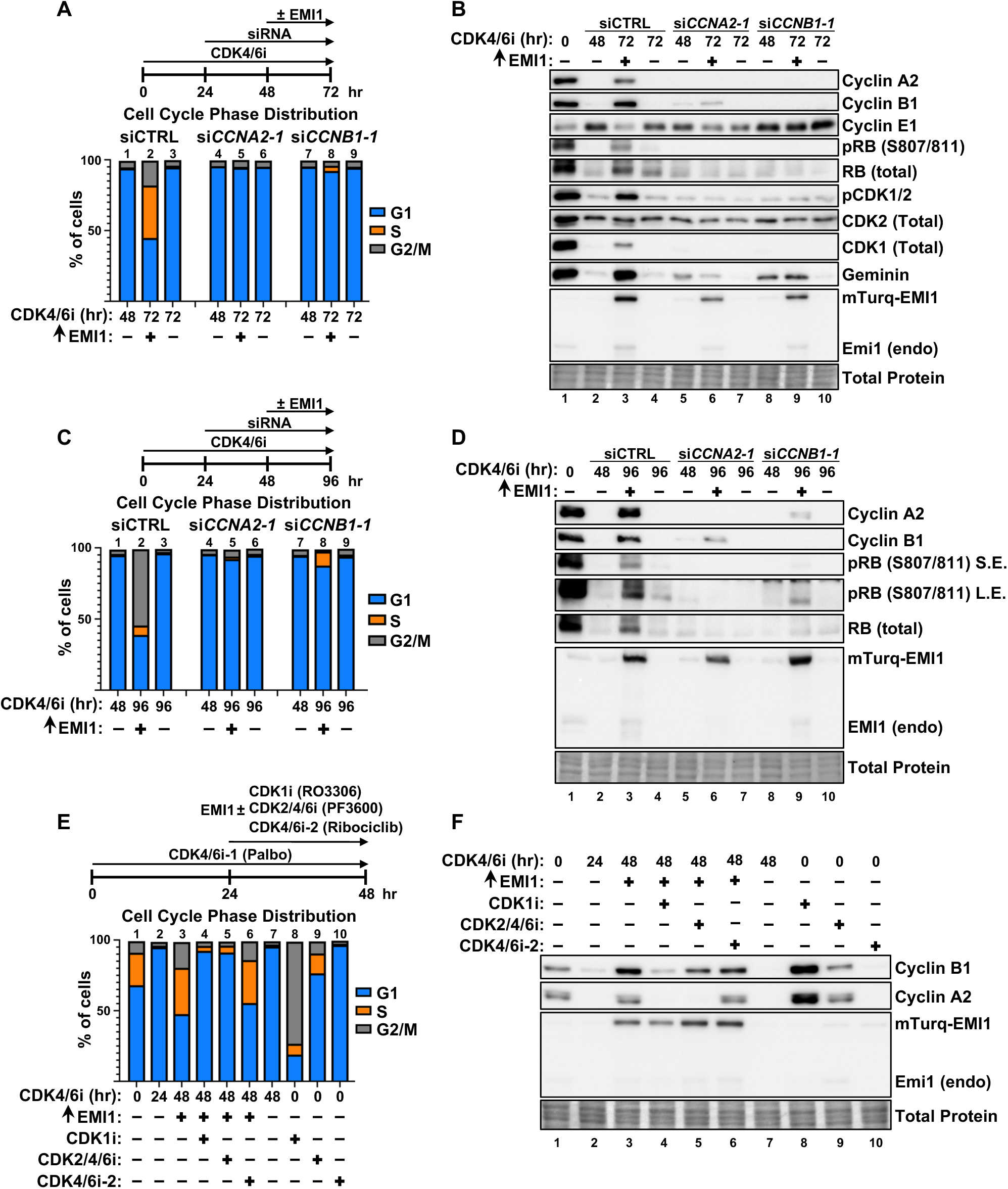
Cyclin/CDK requirements for escape from arrest. **(A)** Cell cycle distribution of RPE1 cells treated with 1 µM CDK4/6i for 24 hours followed by transfection with either control siRNA (“CTRL”) or siRNA targeting either cyclin A (“siCCNA2-1”) or cyclin B (“siCCNB1-1”) for 24 hours prior to 24 hours ± EMI1 expression as indicated (72 hours total; cells remain in CDK4/6i for the entire duration). One sample from each siRNA transfection was collected after 48 hours of CDK4/6i to show that cells were fully arrested at the time of EMI1 induction. **(B)** Immunoblot of lysates from (A). An untreated proliferating sample (lane 1) was included as a control. **(C)** Cell cycle distribution of RPE1 cells treated as in (A) except that EMI1 was induced for the final 48 hours instead of 24 hours (96 hours total; cells remain in CDK4/6i for the entire duration). One sample from each siRNA transfection was collected after 48 hours of CDK4/6i to show that cells were fully arrested at the time of EMI1 induction. **(D)** Immunoblot of lysates from (C). **(E)** Cell cycle distribution of RPE1 cells treated with 1 µM CDK4/6i-1 (Palbociclib) for 24 hours prior to 24 hours of EMI1 expression in the presence of either a CDK1 inhibitor (RO-3306; 5 µM), a CDK2/4/6 inhibitor (PF3600; 25 nM), or a second CDK4/6 inhibitor (Ribociclib; CDK4/6i-2; 5 µM) as indicated (48 hours total; cells remain in first CDK4/6 inhibitor for the entire duration). One sample was collected after 24 hours of CDK4/6i-1 to show that cells were fully arrested at the time of EMI1 induction. RO-3306, PF3600, and Ribociclib were also added to asynchronous cells to show the effects of each single inhibitor. **(F)** Immunoblot of lysates from (E). For cell cycle and immunoblots, one representative example is shown. Panels A-B are representative of 2 biological replicates, panels C-D are representative of 1 biological replicate, and panels E-F are representative of more than 3 biological replicates.

We found no evidence of either CDK T-loop phosphorylation or RB phosphorylation in cyclin A or cyclin B-depleted cells, indicating that all RB-directed CDK activity remained inhibited (Fig. 4*B* and Fig. S6*F*; compare lane 6 to 7 and lane 9 to 10). Notably, the levels of cyclin E protein in arrested cells were equivalent to or higher than in proliferating cells regardless of cyclin A or B suppression (Fig. 4*B*; compare lane 1 to lanes 5 and 8). This result, coupled with the observation that cyclin A or B-depleted cells failed to escape arrest, suggests that cyclin E (at endogenous levels) is insufficient to promote escape in the absence of either cyclin A or cyclin B. Interestingly, we consistently observed that cyclin A depletion did not fully block cyclin B accumulation in EMI1-expressing cells (compare lane 6 to lane 7), whereas cyclin A remained undetectable in cyclin B-depleted cells (compare lane 9 to lane 10) (Fig. 4*B* and Fig S6*F*). These results suggest that cyclins normally associated with later cell cycle phases are more important for S phase entry than G1 cyclins when cells escape from CDK4/6i-induced arrest.

To determine if both cyclins A and B are absolutely required for escape from arrest, we expressed EMI1 in cyclin A or B-depleted cells for a longer time (48 hours). At this timepoint, we observed delayed but detectable S phase entry in cyclin B-depleted cells but found that cyclin A-depleted cells remained completely arrested with no detectable pRB (Fig 4*C-D*). In contrast to expressing EMI1 for 24 hours (Fig 4*B* and S6*F*), cyclin A and pRB were both detectable in cyclin B-depleted cells after 48 hours of EMI1 expression (Fig *4D*; compare lane 9 to lane 10). These data suggest that although cyclin A and B both contribute to escape from arrest, only cyclin A is essential. Cyclin B accumulation accelerates escape from arrest, potentially by promoting cyclin A stabilization or transcription, but ultimately enough cyclin A can accumulate to promote RB phosphorylation and induce S phase entry even in the absence of cyclin B.

The surprising observation that cyclin B, a mitotic cyclin, contributes to escape from arrest prompted us to test the role of specific CDK enzymatic activities. We therefore arrested cells in Palbociclib and then induced EMI1 expression to inhibit the APC/C simultaneously with pharmacological inhibitors targeting either CDK1 or CDK2. We used RO3306 to inhibit CDK1 (43). To inhibit CDK2, we used the CDK2/4/6 inhibitor PF3600, which is more selective for CDK2 than for CDK4/6 at low doses (44, 45). We treated proliferating cells with RO3306 at the indicated dose and observed a strong G2/M arrest, indicating specific CDK1 inhibition (Fig 4*E*; compare column 1 to column 8). Of note, inhibiting CDK1 did not inhibit S phase entry in proliferating cells. Treating proliferating cells with a low dose of PF3600 had minimal effects on overall cell cycle distribution, although we observed a small increase in the percentage of cells in G1 phase and a small decrease in the percentage of cells in S phase, consistent with the expected effect of CDK2 inhibition on S phase entry (Fig 4*E*; compare column 1 to column 9). The fact that PF3600 also inhibits CDK4 and CDK6 is not confounding for our experiments because CDK4 and CDK6 are already inhibited by Palbociclib. However, to confirm that the effects from this inhibitor were due to CDK2 inhibition and not to a “double-hit” of CDK4/6, we also included cells that were first arrested in Palbociclib and then induced to express EMI1 in the presence of a second CDK4/6 inhibitor, Ribociclib. Interestingly, inhibiting either CDK1 or CDK2 prevented escape from Palbociclib-induced arrest when APC/C was also inhibited (Fig 4*E*; compare column 3 to columns 4 and 5), whereas cells treated with Ribociclilb as an extra CDK4/6 inhibitor that does not inhibit CDK2 escaped with similar efficiency as control cells (Fig. 4*E*; compare column 3 to column 6). This result suggests that both CDK1 and CDK2 activities are required for APC/C inhibition to induce bypass of the Palbociclib-induced arrest. Cyclin B accumulated slightly in both CDK1-inhibited and CDK2-inhibited cells (Fig. 4*F*; compare lane 2 to lanes 4 and 5) which phenocopies cyclin A-depleted cells (Fig. 4*B*). In contrast to cyclin B, we did not observe any detectable cyclin A accumulation in either CDK1 or CDK2-inhibited cells (Fig. 4*F*; compare lane 2 to lanes 4 and 5).

From these data, we conclude that APC/C enforces CDK4/6i-induced cell cycle arrest by suppressing CDK1 and CDK2 activity through cyclin A degradation. APC/C-mediated cyclin B degradation also contributes to the arrest, potentially by suppressing the rate of cyclin A accumulation in the absence of APC/C activity, but cyclin B alone is not absolutely required for escape. Given enough time following APC/C inhibition, cyclin A can accumulate to levels sufficient to promote S phase entry even in the absence of cyclin B.

### Cyclin A but not cyclin B is sufficient to promote escape

Having established that both cyclin A and cyclin B contribute to escape from arrest after EMI1 expression, we asked whether expression of either cyclin alone is also *sufficient* to induce escape. Given that inhibiting the APC/C in CDK4/6-inhibited cells leads to cyclin A or B accumulation to levels observed in untreated cells, but not higher levels, we wished to “re-express” these cyclins rather than overexpress them. However, we predicted that at low levels, the APC/C (active in arrested cells) would degrade the ectopic cyclins as rapidly as they are synthesized. To achieve near-normal cyclin expression, we mutated the APC/C degron motifs in cyclin A and cyclin B such that each cyclin can be stably expressed even in cells where the APC/C is active (Fig. 5*A*) (46, 47). We then established new RPE1 cell lines with doxycycline (dox)-inducible expression of either of these variant cyclins.

**Figure 5.**
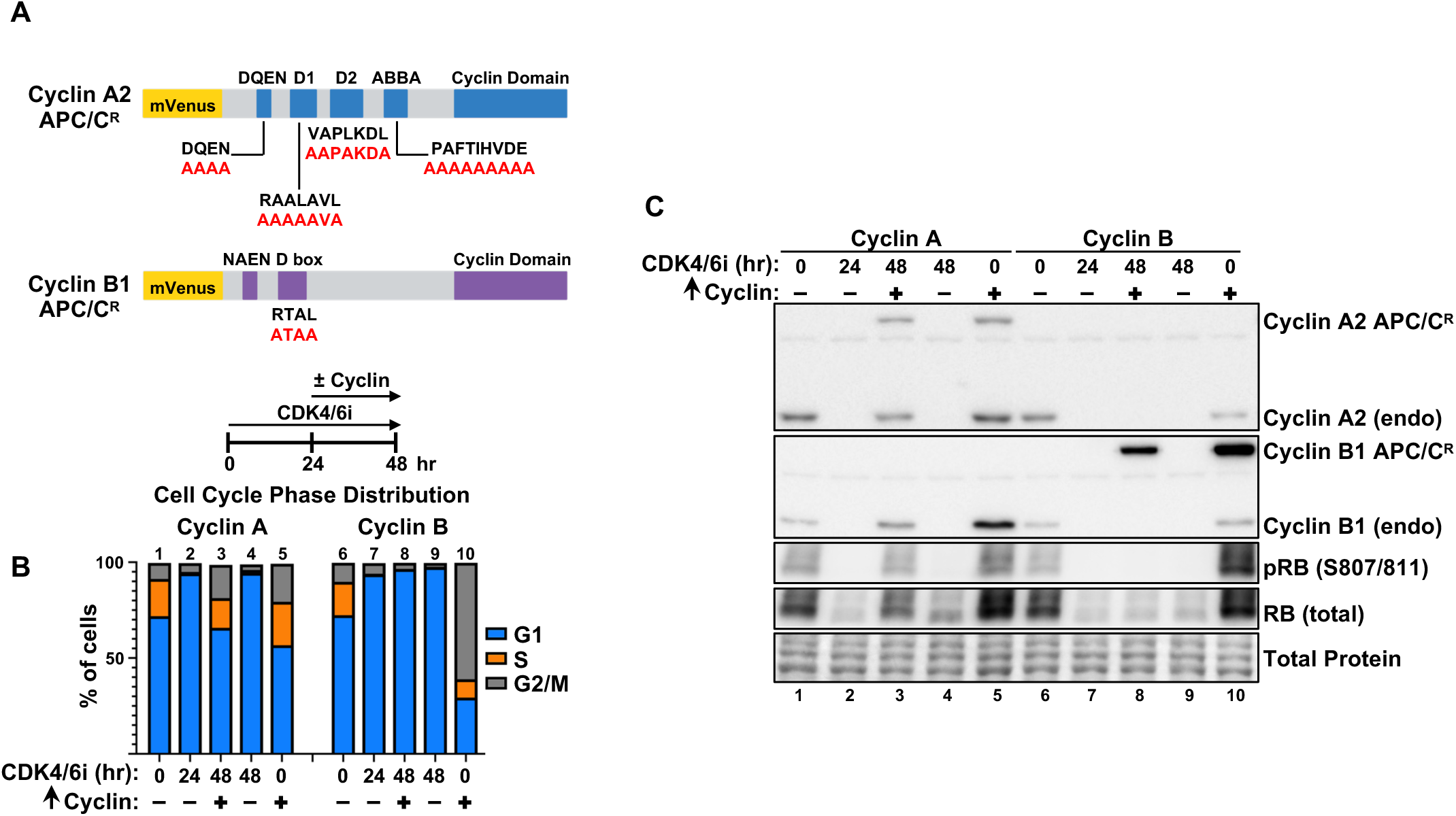
Cyclin A, but not B, expression is sufficient to bypass arrest. **(A)** Schematic of APC/C resistant cyclin A or cyclin B. The APC/C-targeting degron motifs were altered as indicated in red. **(B)** Cell cycle distribution of RPE1 cells treated with 1.25 µM CDK4/6i for 48 hours ± APC/C-resistant cyclin A or B expression during the final 24 hours as indicated. One sample was collected after 24 hours of CDK4/6i to show that cells were fully arrested at the time of cyclin induction. APC/C resistant cyclin A or B were also expressed in asynchronously proliferating cells for 24 hours to confirm that the proteins induce expected cell cycle perturbations (column 1 compared to 5 and column 6 compared to 10). **(C)** Immunoblot of lysates from (B). An untreated proliferating sample from each cell line (lane 1 and lane 6) was included as a control. For cell cycle and immunoblots, one representative example is shown. Panel B is representative of more than 3 biological replicates and panel C is representative of 2 biological replicates.

We found that expression of cyclin A at endogenous levels in arrested cells was sufficient to induce escape from arrest (Fig. 5*B-C*; compare lanes 1-3 in 5*C*). In contrast, cyclin B expression in arrested cells was insufficient to induce either RB phosphorylation or S phase entry, even when cyclin B was overexpressed (Fig. 5*B-C*; compare lanes 6-8 of 5*C*). Expression of stable cyclin B in proliferating cells caused G2/M accumulation because cells unable to degrade cyclin B cannot exit mitosis (48, 49), confirming that the cyclin B variant is active (Fig. 5*B-C*; compare column 6 to column 10 of 5*B*). We conclude that although both cyclin A and cyclin B contribute to escape from CDK4/6i-induced arrest following APC/C inhibition, only direct expression of cyclin A, but not cyclin B, is sufficient to induce escape into S phase.

### Severe underlicensing and proliferation defects in cells bypassing arrest

We recently reported that CDK4/6 inhibition results in the downregulation of several components of the replisome, including members of the minichromosome maintenance (MCM) complex that license DNA replication origins during G1 phase and then form the core of the CMG (Cdc45, MCM2-7, GINS) replicative helicase during S phase (50). In that study, we demonstrated that MCM downregulation during prolonged arrest resulted in underlicensed S phase when cells were allowed to re-enter the cell cycle following drug washout. Reduced levels of chromatin-bound MCM caused inefficient DNA replication, replication stress, and ultimately long-term cell cycle withdrawal (50). Given that in this current study cells escaping arrest are still in the presence of the CDK4/6 inhibitor, we hypothesized that escaping cells are similarly underlicensed. To test for underlicensing, we used a previously published flow cytometry assay to quantify the amount of chromatin-bound MCM in single cells (51, 52). Prior to harvesting cells, we pulse-labeled samples with the nucleotide analog EdU for 1 hour to label S phase cells. During sample collection, we detergent-extracted all soluble proteins such that only chromatin-bound MCM remained. We then stained cells for chromatin-bound MCM2 (a marker of the entire MCM2-7 complex), for EdU, and for DNA content using DAPI. Because cells cannot load any MCM onto DNA once S phase has started, we specifically analyzed the “Early S phase” cells for chromatin-bound MCM content, which represents the maximum amount of MCM cells have available to replicate their DNA (53, 54). We define “Early S phase” cells as those that still have 2C DNA content but are EdU-positive (Fig. 6*A*). Using this assay, we found that cells escaping arrest after EMI1 expression start S phase with less than 15% of the normal amount of chromatin-bound MCM, a level that is severely underlicensed (Fig. 6*B*).

**Figure 6.**
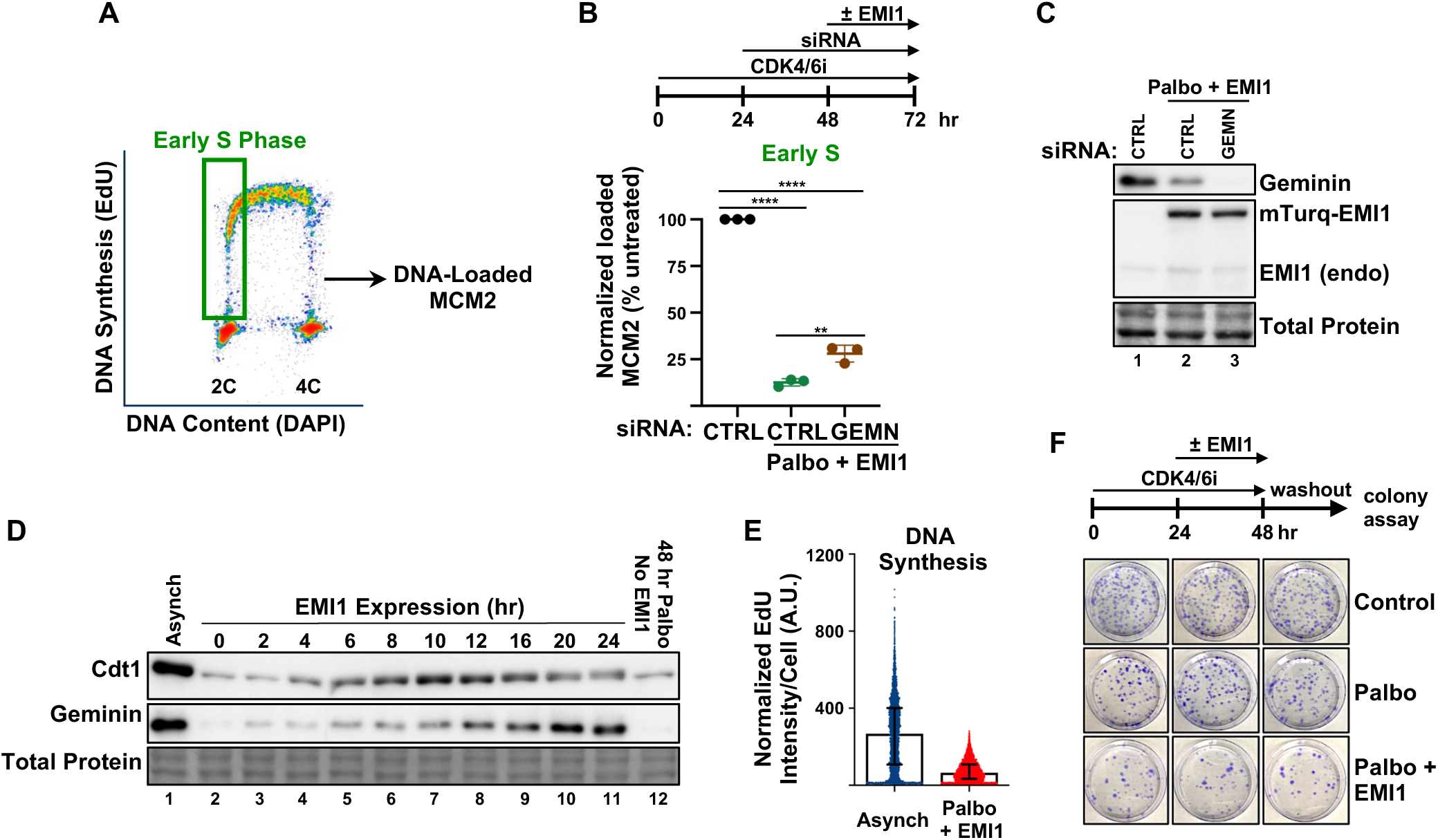
Severe underlicensing and proliferation defects in cells bypassing arrest. **(A)** Gating strategy used to quantify DNA-loaded MCM2 in early S phase cells. The DNA-loaded MCM2 was analyzed from cells that were EdU+ but still had a G1 (2C) DNA content. **(B)** Normalized DNA-loaded MCM2 in early S phase cells. RPE1 cells were treated with 1 µM CDK4/6i for 24 hours followed by transfection with either control siRNA (“CTRL”) or siRNA targeting geminin (“GEMN”) for 24 hours prior to 24 hours of EMI1 expression (72 hours total; cells remain in CDK4/6i for the entire duration). Data displayed are the normalized median DNA-loaded MCM2 intensities of early S phase cells as a percentage of the amount of DNA-loaded MCM2 in untreated early S phase cells. Error bars display the mean ± SD of n = 3 biological replicates. Significance determined by one-way ANOVA followed by Tukey’s multiple comparisons test (**, P ≤ 0.01; ****, P ≤ 0.0001). See materials and methods for normalization method. **(C)** Immunoblot of lysates from (B). **(D)** Immunoblot of lysates from RPE1 cells treated with 1 µM CDK4/6i for 24 hours and then induced to express EMI1 for 0-24 hours. An untreated proliferating sample (“Asynch”; lane 1) was included as a control. Lysates are the same as in figure 1E. **(E)** Normalized EdU intensity per cell of S phase cells from either asynchronously growing RPE1 cells or of S phase cells from RPE1 cells treated with 1 µM CDK4/6i for 48 hours with EMI1 expression during the final 24 hours. Error bars display the median intensity with the interquartile range. See materials and methods for normalization method. **(F)** RPE1 cells were treated with 1 µM CDK4/6i for 48 hours ± EMI1 expression during the final 24 hours as indicated and CDK4/6i and doxycycline were washed out and cells were re-plated at low density and assessed for colony-forming ability. For immunoblots, EdU incorporation, and colony assays, one representative example is shown. Panels C, E, and F are representative of more than 3 biological replicates. Panel D is representative of 2 biological replicates.

Interestingly, geminin – an inhibitor of the essential MCM loading factor CDT1 – is a substrate of the APC/C. During a normal G1 phase while MCM is being loaded, geminin is suppressed by APC/C activity so that cells can properly license origins for replication. Only after cells inactivate the APC/C near the onset of S phase and begin DNA replication does geminin accumulate and inhibit re-licensing and re-replication throughout S/G2/M phases (42, 55). However, we considered that APC/C inhibition in Palbociclib-arrested cells may cause premature geminin accumulation at the same time MCM is being loaded in escaping cells. Indeed, geminin increased within the first 2 hours of APC/C inhibition, with a further increase at 6-8 hours – around the time when cells typically begin to enter S phase (Fig. 6*D*; compare lanes 2-3 and compare lane 4 to lanes 5-6). We predicted that this inappropriate geminin expression was at least partially responsible for underlicensing in escaping cells. In agreement with this prediction, preventing geminin accumulation by RNAi partially rescued MCM loading (Fig. 6*B-C*). We note that while geminin-depleted cells entered S phase with more chromatin-bound MCM than control escaping cells, they were still severely underlicensed compared to untreated control cells.

It is well-established that reduced origin licensing causes replication stress and DNA damage, contributes to proliferation defects, and promotes tumorigenesis (56–64). We thus analyzed the downstream consequences for escaping cells that started S phase severely underlicensed. We found that the rate of DNA replication in escaping cells was dramatically reduced compared to S phase in proliferating cells, demonstrated by a substantially lower EdU incorporation rate per cell (Fig. 6*E*). We next determined the consequences of underlicensed DNA replication in escaping cells for long-term proliferation. We first arrested cells in Palbociclib for 24 hours, followed by an additional 24 hours in the presence or absence of EMI1 expression to inhibit APC/C; at this timepoint, ~50% of cells will have escaped G1 and undergone DNA replication (Fig. 1*D*). We then washed out the dox/inhibitors and re-plated the cells at low density to assess their ability to form colonies. Palbociclib treatment alone for 48 hours prior to re-plating caused only a slight reduction in colony formation compared to untreated control cells, consistent with our previous findings (Fig. 6*F*) (50). However, we found that APC/C inhibition during the final 24 hours in Palbociclib severely impaired colony formation (Fig. 6*F*). We conclude that cells escaping arrest start S phase severely underlicensed, at least in part due to premature geminin accumulation. This underlicensed S phase leads to inefficient DNA replication and long-term proliferation defects.

### APC/C activity is required for CDK4/6 inhibitors to maintain arrest in T47D breast cancer cells

Thus far, we have demonstrated that APC/C inhibition in Palbociclib-arrested RPE1 cells bypasses arrest by causing inappropriate accumulation of cyclins A and B, RB inactivation, and E2F-dependent transcription; arrest bypass correlates with long-term proliferation defects. Given the clinical use of CDK4/6 inhibitors in patients with estrogen receptor (ER)-positive/Her2-negative breast cancer (65), we tested if APC/C activity is also essential to maintain arrest in breast cancer cells. We established a T47D breast cancer cell line with doxycycline (dox)-inducible EMI1, analogous to the RPE1 cell line described above. As we observed in RPE1 cells, APC/C inhibition in Palbociclib-arrested T47D cells caused the reappearance of cyclins A and B, T-loop phosphorylated CDK, and phosphorylated RB (Fig. 7*B*; compare lanes 2-3). These changes were sufficient to induce re-expression of E2F-dependent genes and promote escape from arrest (Fig. 7*A,C*). Using pharmacological inhibitors as before, we confirmed that both CDK1 and CDK2 activities are required for escape from arrest, and that cyclin A accumulation depends on both CDK1 and CDK2 activities (Fig. 7*D-E*; compare lane 3 to lanes 4-5 of 7*E*). Analysis of escaping cells revealed reduced EdU incorporation relative to asynchronously growing cells (Fig. 7*F*). We conclude that APC/C inhibition in Palbociclib-arrested T47D breast cancer cells promotes escape from arrest by a mechanism analogous to that described in RPE1 cells above.

**Figure 7.**
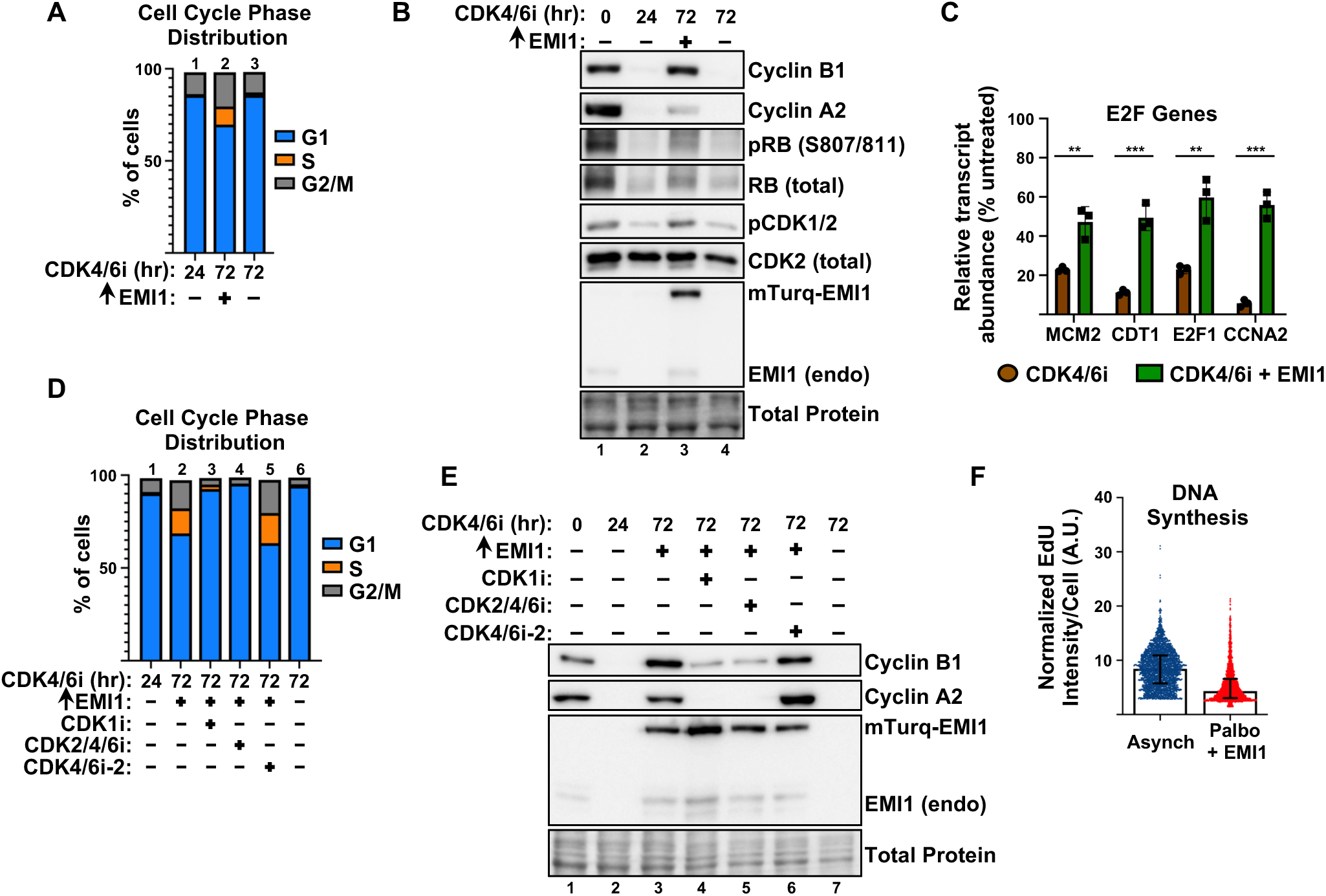
APC/C activity is essential to maintain CDK4/6 inhibitor-induced G1 arrest in T47D breast cancer cells. **(A)** Cell cycle distribution of T47D cells treated with 1 µM CDK4/6i for 72 hours ± EMI1 expression during the final 48 hours as indicated. One sample was collected after 24 hours of CDK4/6i to show that cells were fully arrested at the time of EMI1 induction. **(B)** Immunoblot of lysates from (A). An untreated proliferating sample (lane 1) was included as a control. **(C)** Relative mRNA abundance of E2F-regulated transcripts from T47D cells treated with 1 µM CDK4/6i for 72 hours ± EMI1 expression during the final 48 hours as indicated. Transcript abundances are normalized to a proliferating control sample. Error bars display the mean ± SD of n = 3 biological replicates. Significance determined by unpaired t test (**, P ≤ 0.01; ***, P ≤ 0.001; ****, P ≤ 0.0001). **(E)** Cell cycle distribution of T47D cells treated with 1 µM CDK4/6i-1 (Palbociclib) for 24 hours prior to 48 hours of EMI1 expression in the presence of either a CDK1 inhibitor (RO-3306; 5 µM), a CDK2/4/6 inhibitor (PF3600; 25 nM), or a second CDK4/6 inhibitor (Ribociclib; CDK4/6i-2; 5 µM) as indicated (72 hours total; cells remain in first CDK4/6 inhibitor for the entire duration). One sample was collected after 24 hours of CDK4/6i-1 to show that cells were fully arrested at the time of EMI1 induction. **(F)** Immunoblot of lysates from (D). **(G)** Normalized EdU intensity per cell of S phase cells from either asynchronously growing T47D cells or of S phase cells from T47D cells treated with 1 µM CDK4/6i for 72 hours with EMI1 expression during the final 48 hours. Error bars display the median intensity with the interquartile range. See materials and methods for normalization method. For cell cycle, immunoblots, and EdU incorporation, one representative example is shown. Panels A-B and D-E are representative of 2 biological replicates and panel F is representative of 3 biological replicates.

### High *FBXO5* expression predicts poor prognosis in cancer patients

Our findings that APC/C inactivation impairs cell cycle arrest from CDK4/6 inhibition but causes underlicensing, which is associated with replication stress and genome instability (56–64), suggest that reduced APC/C activity in cancers may promote genome instability, leading to worse patient prognosis. We analyzed publicly available tumor data from The Cancer Genome Atlas (TCGA) Pan-Cancer data set and found that high *FBXO5* expression (EMI1) indeed correlates with reduced overall survival both in pan-cancer (Fig. 8*A*) and breast tumor datasets (Fig 8*B*). These data support the hypothesis that differences in APC/C activity may drive genome instability and cancer progression and be a useful biomarker in predicting patient response to cytostatic therapeutic treatment.

**Figure 8.**
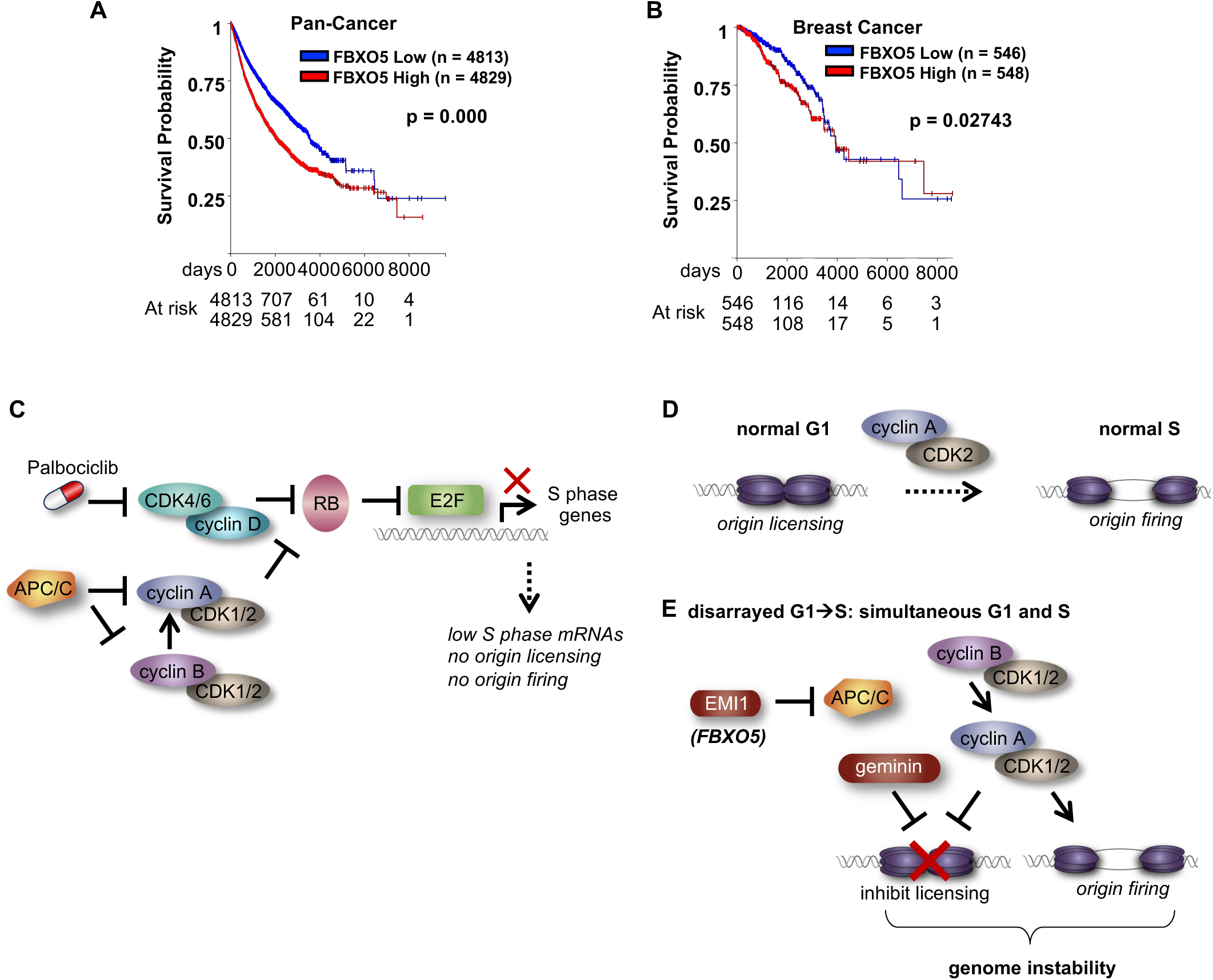
Increased *FBXO5* expression is associated with worse overall survival in cancer patients. Kaplan-Meier overall survival curves from TCGA Pan-Cancer (PANCAN) dataset based on *FBXO5* mRNA expression in **(A)** All primary tumors or **(B)** All primary breast tumors. Samples were separated into high or low *FBXO5* expression based on the median. P value was calculated using the log-rank test. Plots generated using UCSC Xena Browser. **(C)** Model displaying how APC/C activity enforces Palbociclib-induced cell cycle arrest; APC/C-mediated degradation of both cyclin A and cyclin B keeps CDK activity low, which allows RB to repress E2F-dependent transcription and prevent S phase entry. When APC/C activity is inhibited in arrested cells, cyclin B stabilization promotes cyclin A accumulation, which activates CDK and promotes RB phosphorylation and S phase entry. **(D)** DNA replication origin licensing and firing during a normal G1/S transition. Origins are licensed during G1 but remain inactive. Cyclin A/CDK2 activation both promotes origin firing in early S phase and prevents further origin licensing. **(E)** Disarrayed G1/S progression. APC/C inhibition in arrested cells leads to premature stabilization of licensing inhibitors (geminin, cyclin A), preventing licensing during G1. The resulting S phase is underlicensed and results in genome instability and proliferation defects.

## Discussion

APC/C^CDH1^ activity prevents premature G1 progression and S phase entry in proliferating cells (15–23). When mRNA levels are abundant, regulated ubiquitin-dependent degradation is likely to be the predominant control over protein abundance. In contrast, once cells arrest, the transcript levels of cell cycle-promoting genes are much lower (Fig. 2*C,F*, Fig. 7*C*). At the outset of this study it was not clear if repressed gene expression in arrested cells is sufficient to maintain arrest or if continued APC^CDH1^-dependent proteolysis is equally important. Our results demonstrate that, despite highly-repressed E2F-dependent gene expression, APC/C^CDH1^-mediated degradation of cyclins A and B remains essential to keep their protein levels low, avoid RB phosphorylation, and maintain arrest (Fig. 8*C*). Our findings complement several studies in model vertebrate systems showing that loss of APC/C activity in quiescent cells can induce cell cycle re-entry, although this effect may be context- and cell type-dependent (66–69). To our knowledge, our study is the first to demonstrate that APC/C^CDH1^ activity is required to maintain drug-induced arrest.

Importantly, and distinct from studies of normally proliferating cells or tissues, we show that APC/C plays a critical role in *maintaining* the arrest induced by pharmacological CDK4/6 inhibitors. Considerable evidence has accumulated that the cell cycle arrest induced by these inhibitors is unique relative to physiological arrests. Unlike most cells arrested by growth-factor withdrawal or contact inhibition, CDK4/6i-treated cells continue to increase in cell size, rapidly acquire molecular features associated with senescent cells, and suffer genome damage when re-entering the cell cycle (50, 70–75). It is worth noting that EMI1 expression in drug-arrested cells induced a particularly robust escape from arrest compared to escape from contact inhibition or serum withdrawal (Fig. S3). These differences could be related to the relative synchrony of the different arrested populations, but they could also reflect drug-arrested cells poised on the brink of G1 progression because they experience few other growth inhibitory signals. Future studies of distinct cell cycle arrest states may shed light on these differences.

Our data provide direct evidence that cyclin A is the key APC/C^CDH1^ substrate that must be degraded to prevent premature proliferation. Surprisingly, we also discovered that cyclin B stabilization in arrested cells substantially accelerates S phase entry. However, unlike cyclin A, which we demonstrate is sufficient to bypass drug arrest, cyclin B is neither essential nor sufficient (Fig. 4*C-D*, 5*B-C*). Interestingly, cyclin B-depleted cells showed delayed cyclin A accumulation (Fig. 4*D* and 8*C*). Despite primarily being associated with CDK1 activity driving mitosis, several reports indicate a potential role for cyclin B at the G1/S transition, at least in some contexts (67, 76, 77). However, directly expressing cyclin B in arrested cells did not lead to any RB phosphorylation, cyclin A mRNA or protein accumulation, or any S phase entry (Figure 5*B-C*). We postulate that at least one additional APC/C^CDH1^ substrate cooperates with cyclin B to drive cyclin A accumulation and S phase entry. One intriguing candidate for a cooperating substrate is the PLK1 kinase. PLK1 was strongly downregulated in arrested cells and reaccumulated following APC/C inhibition (Supp table 1). In proliferating cells, Cyclin B/CDK1 phosphorylates the FOXM1 transcription factor to prime it for phosphorylation by PLK1, which activates its transcriptional activity, and FOXM1 binds the *CCNA2* (cyclin A) promoter (78–81). Co-stabilizing cyclin B and PLK1 by APC/C inhibition may thus induce FOXM1-dependent transcription of *CCNA2*, accelerating the rate of escape from arrest.

We were surprised to discover that cyclin B acts before cyclin A to drive S phase entry in drug-arrested cells. G1 progression in proliferating cells is controlled by Cyclin D CDK4/6 and Cyclin E/CDK2 followed by cyclin A activity beginning from very early S (with CDK2) through G2 phase (with CDK1); cyclin B/CDK1 then acts in G2 and M and is not typically associated with S phase entry. This non-canonical order of action - cyclin B before A, and both mitotic cyclins driving S phase independently of cyclin D and (apparently) cyclin E – is the reverse of a typical cell cycle. Thus, escape from drug-induced arrest is not by simply reactivating the normal G1-S progression program but rather by following a sequence of molecular events not associated with normal cell cycle progression. Other variant sequences of molecular cell cycle events have also been recently described in perturbed cell cycles (82–84).

We were also surprised that cells arrested in Palbociclib display elevated cyclin E relative to proliferating cells yet remained fully arrested (Fig. 1*E*). Cyclin E was equally elevated even in cyclin A and cyclin B-depleted cells which remained arrested following APC/C inhibition (Fig. 4*A-B*). Thus, cyclin E levels showed no correlation with escape or arrest outcomes. However, we and others have shown that very high cyclin E overexpression *can* drive premature S phase in proliferating cells (51, 85–88). Moreover, high cyclin E protein (89) and high *CCNE1* mRNA (90, 91) correlate with reduced CDK4/6i sensitivity. Models of acquired drug resistance report cyclin E overexpression or amplification in the resulting resistant population (92–95). However, in the context of CDK4/6 inhibitor-induced arrest, endogenous cyclin E is clearly insufficient to drive escape from arrest. What keeps cyclin E/CDK2 from phosphorylating RB and driving S phase entry in these cells? The essential activating T-loop phosphorylations on CDK2 at Thr160 and CDK1 at Thr161 (96, 97) are abolished by Palbociclib treatment (Fig. 1*E*). The CDK-activating kinase (CAK) that phosphorylates CDK T-loops is thought to be constitutively active (98). However, the phosphatase that antagonizes this phosphorylation (KAP) preferentially dephosphorylates monomeric CDK2, and binding of cyclin A protects it from dephosphorylation (99). Moreover, CAK can preferentially phosphorylate cyclin/CDK complexes over monomeric CDK, particularly CDK1 (100). We thus speculate that premature cyclin A accumulation leads to CDK re-activation at least in part by facilitating activating CDK phosphorylation.

We discovered that, despite the rapid S phase entry in cells escaping from drug-induced arrest, DNA replication and proliferation were profoundly impaired (Fig. 6*E-F*). The replication defect is characterized by severe origin underlicensing (Fig 6*B*). We propose that the cause of underlicensing is a combination of reduced MCM and CDT1 levels (Fig. 6*D*) with the simultaneous accumulation of two APC/C substrates that are both potent licensing inhibitors, geminin and cyclin A (42, 101). Instead of the normal order of unimpeded origin licensing in G1 followed by normal S phase in which origin licensing is inhibited to avoid toxic re-replication while origins are firing (reviewed in (53)), we presume that escaping cells are licensing origins, firing origins, and inhibiting licensing at the same time (Fig 8*D-E*). APC/C activity ensures origin licensing and firing are temporally separated, but EMI-induced APC/C inhibition removes this strict separation. This model is consistent with the documented role of APC/C^CDH1^ in maintaining genome integrity (17–19, 21) and the documented replication stress and genome damage caused by underlicensing (reviewed in (102)).

CDK4/6 inhibitors have revolutionized the treatment of metastatic estrogen receptor (ER)-positive/Her2-negative breast cancer (65, 103). However, both intrinsic and acquired resistance to CDK4/6 inhibitor treatment continues to limit their therapeutic efficacy (104). Our data suggest that tumors with naturally reduced APC/C^CDH1^ activity (e.g. elevated *FBXO5* (EMI1) expression) may exhibit intrinsic resistance and continue to divide. Supporting this idea, knockdown of CDH1 *prior* to CDK4/6 inhibitor treatment in T47D breast cancer cells reduced the percentage of cells establishing arrest (24). One natural extension of this observation is that markers of reduced APC/C^CDH1^ activity may be a useful biomarker to predict treatment response. On the other hand, CDK4/6 inhibitors could force tumor cells with low APC/C^CDH1^ activity into a self-destructive underlicensed cell cycle sensitive to replication-targeting treatments. Non-tumor cells with normal APC/C activity would be protected because they would remain successfully arrested in G1 phase and thus not undergoing DNA replication. A similar strategy exploiting elevated replication stress following release from extended Palbociclib treatment has recently been shown to be effective in promoting stable cell cycle exit (50). Future studies will determine how our findings that APC/C^CDH1^ is essential to maintain arrest induced by CDK4/6 inhibition can be leveraged in the clinic to either predict patient response or enhance therapeutic efficacy.

## Materials and Methods

### Cell culture

All cell lines utilized were verified to be mycoplasma negative. RPE1-hTERT and HEK293T cells were cultured in DMEM (Sigma-Aldrich Chemistry) supplemented with 10% FBS (Avantor Seradigm), 2 mM L-glutamine (Gibco), and 1x pen/strep (Gibco). T47D cells were cultured in phenol-red free RPMI-1640 (Gibco) supplemented with 10% FBS, 2 mM L-glutamine, 10 µg/mL human insulin (Gibco), and 1x pen/strep. Cells were incubated at 37°C in 5% CO_2_ and were passaged with 1x trypsin (Sigma-Aldrich Chemistry) before reaching confluency.

### Inhibitors/Chemicals

The following inhibitors/chemicals were used in this study: PD0332991 (Palbociclib, CDK4/6i, Selleckchem, Cat # S1116), LEE011 (Ribociclib, CDK4/6i, Selleckchem, Cat # S7440), RO-3306 (CDK1i, Selleckchem, Cat # S7747), PF06873600 (CDK2/4/6i, Selleckchem, Cat # S8816), HLM006474 (E2Fi, Selleckchem, Cat # S8963), Doxycycline (CalBiochem, Cat # 32485).

Concentrations used are as follows unless otherwise indicated: Palbociclib, 1 µM (except for figure 5 where 1.25 µM was used); Ribociclib, 5 µM; RO-3306, 5 µM, PF06873600, 25 nM; HLM006474, 20 µM (higher concentrations of the E2F inhibitor may have produced stronger phenotypes but also induced apparent toxicity); Doxycycline, 1 µg/mL.

### Cell line generation

The RPE1-hTERT cells containing inducible mTurq-EMI1 were a gift from R. Medema and have been published previously (25)

All new stable cell lines generated in this study were made by cloning the cDNA of interest into lentiviral expression plasmids followed by viral transduction and drug selection of cells. The pCW-puro_mTurq-EMI1-WT plasmid was a gift from R. Medema and has been published previously (25). The pTRIPZ-puro_2xFLAG-2xSTREP-Emi1-WT and pTRIPZ-puro_2xFLAG-2xSTREP-Emi1-ΔF-box plasmids were gifts from M. Pagano and have been published previously (34).

To make plasmid encoding inducible E2F2: The FUW-tetO-hOKMS plasmid (plasmid #51543; Addgene) was cut with EcoR1 to linearize it and remove the original OKMS insert. The pCMVHA E2F2 plasmid (plasmid #24226; Addgene) was used as a template for amplifying the HA-tagged E2F2 gene using conventional PCR. Primers used were: Vector-E2F2.FOR: 5’-<u>gatccagcctccgcggcccc</u>gaattcatggcatacccatacgatgtt-3’ and Vector-E2F2.REV: 3’-gaacccctggacaactaattaactcttaa<u>gctatagttcgaatagctatt</u>-5’ with overlapping regions (underlined) with the linearized FUW-tetO plasmid. The amplified HA-E2F2 fragment was assembled into the linearized FUW-tetO plasmid by Gibson Assembly (New England Biolabs) following the manufacturer’s protocol. Successful cloning was confirmed by Sanger sequencing. The generated plasmid was designated as FUW-tetO-HA-E2F2.

To make plasmids encoding inducible APC/C resistant cyclin A or cyclin B: cDNAs encoding either APC/C resistant (APC/C^R^) mVenus-cyclin A2 or mVenus-cyclin B1 were cloned into pINDUCER20 (plasmid #44012; Addgene) by replacing the gateway cassette sequence and were synthesized by Twist Biosciences. The mutations in the APC/C degron motifs of cyclin A were designed as described previously (46). The D box of cyclin B1 was mutated from 42-RTAL-45 to 42-ATAA-45.

To generate cell lines: lentiviral expression plasmids were co-transfected with ΔNRF and VSVG (gifts from J. bear) virus packaging plasmids into HEK293T cells using 50 µg/mL polyethylenimine (PEI)-Max (Sigma-Aldrich Chemistry). Two days later, virus-containing supernatant was collected and used to transduce cells (either RPE1-hTERT or T47D; to make the cells expressing inducible E2F2: RPE1-hTERT cells constitutively expressing rtTA (reverse tetracycline Trans Activator) were used as target cells) in the presence of 8 µg/mL polybrene (Millipore) for at least 6 hours (maximum 24 hours). In some cases, a second round of viral transduction was performed. Cells that successfully integrated the transduced constructs were selected by treating cells with either puromycin (Sigma-Aldrich Chemistry) or geneticin (Gibco) until mock-infected cells (incubated with polybrene-containing media with no virus) treated with the same concentration of drug were completely dead. All cell lines generated are polyclonal except for the RPE1-hTERT cells expressing inducible E2F2 and the RPE1-hTERT cells expressing inducible APC/C^R^ cyclin A. For the RPE1-hTERT cells expressing inducible E2F2: cells were plated for clonal selection following viral transduction (no drug selection) and clones were screened for expression of the construct of interest. For the RPE1-hTERT cells expressing inducible APC/C^R^ cyclin A: cells were plated for clonal selection following viral transduction and drug selection and clones were screened for expression of the construct of interest.

### Immunoblotting

Following harvesting of samples by trypsinization: cell pellets were lysed in cold CSK lysis buffer (300 mM sucrose, 300 mM NaCl, 3 mM MgCl_2_, and 10 mM PIPES pH 7.0) supplemented with 0.5% triton X-100 (Sigma-Aldrich Chemistry), protease inhibitors (0.1 mM AEBSF, 1 µg/mL pepstatin A, 1 µg/mL leupeptin, and 1 µg/mL aprotinin), and phosphatase inhibitors (10 µg/mL phosvitin, 1 mM β-glycerol phosphate, and 1 mM Na-orthovanadate) and incubated on ice for 15 minutes. Lysates were clarified by centrifugation at 16,000*g* for 10 minutes at 4°C. Bradford assay (Biorad) was performed on the clarified lysates to determine the protein concentrations and all samples were diluted in lysis buffer and SDS loading buffer (final concentrations: 1% SDS, 2.5% 2-mercaptoethanol, 0.1% bromophenol blue, 50 mM Tris pH 6.8, and 10% glycerol) such that the final concentration of protein was 1 µg/uL. Samples were boiled for 5 minutes before separating 15-20 µg of protein on SDS-PAGE gels. Proteins were transferred to polyvinylidene difluoride membranes (PVDF; Thermo Fisher Scientific) and membranes were blocked in 5% milk diluted in TBS with 0.1% Tween 20 (TBS-T) for 1 hour at room temperature, shaking. Following blocking, membranes were incubated in primary antibody diluted in either 5% milk (TBST) or 5% bovine serum albumin (BSA; Thermo Fischer Scientific) (TBST) either overnight in cold room or for 1 hour at room temperature, shaking. Membranes were washed 3x in TBST for 10 minutes/each followed by incubation in HRP-conjugated secondary antibody diluted in 5% milk (TBST) for 1 hour at room temperature, shaking. Membranes were washed 3x in TBST for 10 minutes/each and imaged with ECL Prime (Amersham) using ChemiDoc MP imaging system (BioRad). Total protein was visualized using Ponceau S stain (Sigma-Aldrich).

The following antibodies were used in this study: *Cyclin A2* (1:2000; mouse; 4656; Cell Signaling Technology; milk), *pRB S807/811* (1:1000; rabbit; 9308; Cell Signaling Technology; BSA), *Total RB* (1:3000; mouse; 554136; BD Biosciences; milk), *FBXO5 (Emi1)* (1:500; mouse; 37-660; Thermo Fisher Scientific; milk), *Cyclin E1* (1:2000; mouse; 4129; Cell Signaling Technology; milk), *pCDK2 T160* (1:1000; rabbit; 2561; Cell Signaling Technology; BSA), *Total CDK2* (1:1000; rabbit; 2546; Cell Signaling Technology; BSA), *E2F1* (1:1000; rabbit; 3742; Cell Signaling Technology; BSA), *E2F2* (1:1000; rabbit; A3297; Abclonal; milk), *E2F3* (1:1000; mouse; sc-56665; Santa Cruz Biotechnology; milk), *Skp2* (1:1000; rabbit; 4313; Cell Signaling Technology; milk), *p27* (1:1000; rabbit; 3686; Cell Signaling Technology; BSA), *Cyclin B1* (1:1000; mouse; sc-245; Santa Cruz Biotechnology; milk), *Total CDK1* (1:10,000; rabbit; A303-664A; Bethyl Laboratories; milk), *Geminin* (1:3000; rabbit; sc-13015; Santa Cruz Biotechnology; milk), *Cdt1* (1:1000, rabbit, 8064, Cell Signaling Technology, milk), donkey-anti-mouse-HRP (1:10,000; 715-035-150; Jackson ImmunoResearch), and donkey-anti-rabbit-HRP (1:10,000; 711-035-152; Jackson ImmunoResearch).

### qPCR

Total cellular RNA was extracted from cells using the RNeasy plus mini kit (Qiagen, cat # 74134). RNA was reverse transcribed into cDNA according to manufacturer’s instructions using the High-Capacity Reverse Transcription Kit (Thermo Fisher Scientific, cat # 4368814). qPCR was performed using iTaq Universal SYBR Green Supermix (Biorad, cat # 1725121) and analyzed on the QuantStudio 7 system (Thermo Fisher Scientific) using the standard sybergreen protocol. For each gene being analyzed: all samples were pipetted in technical triplicate. Transcript abundances were calculated using the 2^^−Δ^ ^ΔCt^ method by first normalizing the mean Ct value of the gene of interest to the mean Ct value of the housekeeping gene, ACTB, followed by a second normalization to the experimental biological control (untreated asynchronous cells).

The following primer sequences were used: *MCM2,* 5’-GCAGCATGCGCAAGACTTT-3’ (fwd) and 5’-TGTCACCTGCTCTGCCACTAA-3’ (rev); *CDT1*, 5’-GTAGGCGTTTTGAGGAGTGC-3’ (fwd) and 5’-TAATCTGACCTCCTGGTGCC-3’ (rev); *E2F1*, 5’-TCTCCGAGGACACTGACAG-3’ (fwd) and 5’-ATCACCATAACCATCTGCTCTG-3’ (rev); *CCNA2*, 5’-CGCTGGCGGTACTGAAGTC-3’ (fwd) and 5’-GAGGAACGGTGACATGCTCAT-3’ (rev); *FBXO5 (EMI1),* 5’-GTGTCTAAAGTGAGCACAACTTG-3’ (fwd) and 5’-TTCTCTGGTTGAAGCATGAGG-3’ (rev); *ACTB*, 5’-ACCTTCTACAATGAGCTGCG-3’ (fwd) and 5’-CCTGGATAGCAACGTACATGG-3’ (rev)

### Flow cytometry for cell cycle analysis

1 hour prior to cell collection, samples were pulsed with 10 µM EdU (Santa Cruz Biotechnology) to label cells actively synthesizing DNA (S phase cells). Cells were harvested by trypsinization, washed 1x in phosphate buffered saline (PBS), and fixed in PBS + 4% paraformaldehyde (Sigma-Aldrich Chemistry) for 15 minutes at room temperature. 1 mL of PBS + 1% BSA was added to the fixed cells and cells were spun at 1,000*g* for 5 minutes. Supernatant was aspirated and cells were stored in 1 mL of PBS + 1% BSA at 4°C until staining.

To stain cells for flow cytometry analysis: Cells were first permeabilized in 500 uL PBS + 1% BSA + 0.5% Triton X-100 for 15 minutes at room temperature. Cells were spun at 1,000*g* for 5 minutes and the supernatant was aspirated. EdU was labeled with Alexa fluor 647-azide (Life Technology) by incubating cells in EdU labeling solution (1 mM CuSO_4_, 1 µM AF-647, 100 mM ascorbic acid, diluted in PBS) for 30 minutes at room temperature in the dark. Cells were spun at 1,000*g* for 5 minutes and the supernatant was aspirated. DNA content was labeled by incubating cells in 1 µg/mL DAPI (Sigma-Aldrich Chemistry) supplemented with 100 µg/mL RNase (Sigma-Aldrich Chemistry) (diluted in PBS + 1% BSA + 0.5% Triton X-100) for either 1 hour at 37°C or overnight at 4°C (in the dark). Cells were analyzed on an Attune NxT flow cytometer (Thermo Fisher Scientific) and the results were analyzed using FCS Express 7 Research (De Novo Software). Cell cycle distribution was determined by gating on a plot of DAPI vs EdU (see figure S1).

To compare EdU intensity per cell between samples (figure 6 and figure 7): Due to an increase in autofluorescence caused by an increase in cell size in Palbociclib-arrested cells, the data were normalized. To do so: for each sample condition (untreated or Palbociclib-treated), a sample that was treated identically but was not stained for EdU (AF-647) was included. The EdU (AF-647) intensities of S phase cells from the stained samples were divided by the median EdU (AF-647) intensity of the corresponding unstained control sample (ex: AF-647 intensities of an untreated but stained sample were normalized to the median AF-647 intensity of an untreated and unstained sample).

### Chromatin-bound MCM flow cytometry

1 hour prior to cell collection, samples were pulsed with 10 µM EdU to label cells actively synthesizing DNA (S phase cells). Samples were harvested by trypsinization, washed 1x in cold PBS, and the soluble protein was pre-extracted from cells by incubating on ice in cold CSK lysis buffer supplemented with 0.5% triton X-100, protease inhibitors, and phosphatase inhibitors (described above for immunoblotting) for 8 minutes. Cells were then washed in 1 mL of PBS + 1% BSA, fixed in PBS + 4% paraformaldehyde for 15 minutes at room temperature, and then washed again in 1 mL of PBS + 1% BSA and stored at 4°C until staining.

To stain cells for flow cytometry analysis: EdU was labeled with Alexa fluor 647-azide by incubating cells in EdU labeling solution (described above) for 30 minutes at room temperature in the dark. 1 mL of PBS + 1% BSA + 0.1% NP-40 was added to cells, cells were spun at 2,000*g* for 10 minutes and the supernatant was aspirated. Chromatin-bound MCM was labeled by incubating cells in antibody recognizing MCM2 (1:200; 610700; BD biosciences) for 1 hour at 37°C in the dark. 1 mL of PBS + 1% BSA + 0.1% NP-40 was added to cells, cells were spun at 2,000*g* for 10 minutes and the supernatant was aspirated. Cells were then incubated in anti-mouse secondary antibody conjugated to Alexa fluor 488-azide (Life Technology, 1:1000) for 1 hour at 37°C in the dark. 1 mL of PBS + 1% BSA + 0.1% NP-40 was added to cells, cells were spun at 2,000*g* for 10 minutes and the supernatant was aspirated. DNA content was labeled by incubating cells in 1 µg/mL DAPI supplemented with 100 µg/mL RNase (diluted in PBS + 1% BSA + 0.1% NP-40) for either 1 hour at 37°C or overnight at 4°C (in the dark). Cells were analyzed on an Attune NxT flow cytometer and the results were analyzed using FCS Express 7 Research.

Due to an increase in autofluorescence caused by an increase in cell size in Palbociclib-arrested cells, the data were normalized. To do so: for each sample condition (untreated or Palbociclib-treated), a sample that was treated identically but was not stained for MCM2 (AF-488) or EdU (AF-647) was included. The MCM2 (AF-488) intensities of early S phase cells (see figure 6A) from the stained samples were divided by the median MCM2 (AF-488) intensity of the corresponding unstained control sample (ex: AF-488 intensities of an untreated but stained sample were normalized to the median AF-488 intensity of an untreated and unstained sample). The medians of these normalized intensities were then normalized to the experimental biological control (untreated asynchronous cells).

### siRNA transfections

siRNAs were transfected into cells using the Dharmafect 4 transfection reagent (Horizon Discovery) according to manufacturer’s instructions. Briefly, the day before transfection, cells were seeded into 10-cm dishes. Transfection mixtures (siRNA + transfection reagent diluted in OPTI-MEM (Gibco)) were prepared to deliver 500 uL to each dish (10% of the final volume). Transfection mixtures contained siRNA such that the final concentrations added to cells are as listed below. For each dish: 1 uL of Dharmafect 4 was used per 100,000 cells transfected (determined by counting an extra dish seeded at the same density). Cells were incubated with the transfection mixture in a final volume of 5 mL of antibiotic-free DMEM + 10% FBS for 24 hours. After 24 hours, cells were washed 1x in 1x PBS and media was replaced with full growth media and the experiment was completed by treatment with the appropriate inhibitors/chemicals/manipulations.

All siRNAs were synthesized by Life Technology. siRNA sequences and concentrations used in this study are as follows:

*siCTRL (siLuciferase):* 25-100 nM (depending on the concentration of other siRNAs in each experiment), 5’-UCGAAGUACUCAGCGUAAGdTdT-3’

*siRB:* 100 nM, 5’-CGAAAUCAGUGUCCAUAAAdTdT-3’

*siCCNA2-1:* 75 nM, 5’-AAGGAUCUUCCUGUAAAUGAUGAGCdTdT-3’

*siCCNA2-2:* 100 nM, 5’-AACUACAUUGAUAGGUUCCUGdTdT-3’

*siCCNB1-1:* 75 nM, 5’-AAGAAAUGUACCCUCCAGAAAdTdT-3’

*siCCNB1-2:* 100 nM, 5’-AAACUUUCGCCUGAGCCUAUUdTdT-3’

*siE2F1:* 25 nM, 5’-GUCACGCUAUGAGACCUCAdTdT-3’

*siGEMN:* 100 nM, 5’-CUUCCAGCCCUGGGGUUAUdTdT-3’

### MS sample preparation for global, quantitative proteomics

All experimental conditions were plated and analyzed in technical triplicate. Following treatment with Palbociclib ± EMI1 expression (8, 16, or 20 hours of EMI1 expression; 16 hour EMI1 expression data included in Supplemental table 1 but not in figure 3*A*), cells were harvested via trypsinization, washed 3x in 1X PBS, and stored at −80°C until sample preparation. Cell pellets were lysed on ice for 10 minutes in urea lysis buffer (8M urea, 75 mM NaCl, 50 mM Tris pH 8.0, 1 mM EDTA) supplemented with protease inhibitors (0.1 mM AEBSF, 1 µg/mL pepstatin A, 1 µg/mL leupeptin, and 1 µg/mL aprotinin), and phosphatase inhibitors (10 µg/mL phosvitin, 1 mM β-glycerol phosphate, and 1 mM Na-orthovanadate). Lysates were snap frozen in ethanol/dry ice slurry 1x and allowed to thaw on ice. Lysates were clarified by centrifugation at 16,000*g* for 15 minutes at 4°C and supernatants were transferred to new Eppendorf tubes. Protein concentrations were determined using BCA assay and lysates were analyzed by mass spectrometry, as described below.

For each sample, 150ug of protein lysate was reduced with 5mM dithiothreitol (DTT; Pierce) at 37°C for 45 minutes and alkylated with 15mM iodoacetamide (IAA; Pierce) for 45 minutes at room temperature. Samples were then precipitated by adding six volumes of ice-cold acetone and incubating at −20°C overnight. The following day, samples were centrifuged at 15000xg for 15 minutes at 4°C, supernatant was removed, and the pellet was washed with 100ul cold acetone. Pellets were air dried at room temperature for 10 minutes before being resuspended in 100ul 1M urea, 50mM ammonium bicarbonate, pH 8. Samples were then subjected to digestion with LysC (Wako) at 37°C for 2 hours and trypsin (Promega) overnight at 37°C at a 1:50 enzyme:protein ratio. The resulting peptides were acidified to 0.5% trifluoroacetic acid (Pierce) and desalted using Thermo desalting spin columns. Eluates were dried via lyophilization, and peptide concentration was determined via Pierce Quantitative Fluorometric Assay and all samples were normalized to 0.25 ug/ul, spiked with iRT peptides (Biognosys) at a 1:30 ratio, and subjected to LC-MS/MS analysis.

### LC-MS/MS for global, quantitative proteomics

Samples were analyzed in a randomized order by LC-MS/MS using an Ultimate 3000 coupled to an Exploris 480 mass spectrometer (Thermo Scientific). A pooled sample was analyzed intermittently throughout the sample set to assess instrument performance. Samples were injected onto an IonOpticks Aurora series 3 C18 column (75 μm id × 15 cm, 1.6 μm particle size) and separated over a 200 min method. The gradient for separation consisted of 3-41% mobile phase B at a 250 nl/min flow rate, where mobile phase A was 0.1% formic acid (Pierce) in LC-MS grade water (Fisher) and mobile phase B consisted of 0.1% formic acid in 80% acetonitrile (Fisher). The Exploris 480 was operated in product ion scan mode for Data Independent Acquisition (DIA).

A full MS scan (m/z 350-1650) was collected; resolution was set to 120,000 with a maximum injection time of 20 ms and automatic gain control (AGC) target of 300%. Following the full MS scan, a product ion scan was collected (30,000 resolution) and consisted of stepped higher collision dissociation (HCD) set to 25.5, 27, 30; AGC target set to 3000%; maximum injection time set to 55 ms; variable precursor isolation windows from 350-1650 m/z.

### Data analysis for global quantitative proteomics

Raw data files were processed using Spectronaut (v17.1.22.229.55965; Biognosys) and searched against the Uniprot reviewed human database (UP000005640, containing 20,396 entries, downloaded March 2021) and the MaxQuant common contaminants database (246 entries). The following settings were used: enzyme specificity set to trypsin, up to two missed cleavages allowed, cysteine carbamidomethylation set as a fixed modification, methionine oxidation and N-terminal acetylation set as variable modifications. Precision iRT calibration was enabled, single hits were excluded, and a false discovery rate (FDR) of 1% was used to filter all data. Normalization was enabled and row selection set to ‘identified in all runs (complete) and missing values were imputed. Un-paired student’s t-tests were conducted and corrected p-values (q-values) were calculated in Spectronaut. One sample (8 hour timepoint replicate 3) was identified as an outlier and removed for analysis.

### Colony Forming Assay

Following treatment with Palbociclib ± EMI1 expression, cells were washed 3x with 1x PBS. Cells were trypsizined and each sample was re-plated in technical triplicate at a density of 500 cells/10-cm dish. Cells were grown for 12 days, replenishing the media every 3 days. After 12 days: cells were washed 2x in 1x PBS, fixed for 10 minutes in 5 mL methanol, stained with 5 mL of 1% (w/v) crystal violet diluted in 20% EtOH for 20 minutes, rinsed lightly with H_2_O, and allowed to dry prior to taking pictures.

### Patient Survival Analysis

Kaplan-Meier curves were generated using publicly available gene expression data found in The Cancer Genome Atlas (TCGA) Pan-Cancer (PANCAN) dataset. Graphs were generated using the University of California Santa Cruz Xena Browser tool (105). For each plot (pan-cancer or breast tumor) samples were filtered such that only the primary tumors in the dataset were analyzed. *FBXO5* low vs. high tumors were demarcated using the median expression value.

### Statistical Analysis

Bar graphs displaying qPCR data and scatter dot plot displaying chromatin-bound MCM flow cytometry data represent the mean and the error bars indicate the SD. Significance tests for qPCR data were performed using unpaired t test for each gene. Significance tests for chromatin-bound MCM flow cytometry data were performed using ordinary one-way ANOVA followed by Tukey’s multiple comparison test. Significance test for Kaplan-Meier curves was performed using the log-rank test. Number and types of replicates are indicated in the figure legends. Statistical significance values are as follows: n.s., not significant; *, P ≤ 0.05; **, P ≤ 0.01; ***, P ≤ 0.001; ****, P ≤ 0.0001. GraphPad Prism was used for statistical analysis.

### Data Availability

The mass spectrometry proteomics data have been deposited to the ProteomeXchange Consortium via the PRIDE [1] partner repository with the dataset identifier PXD046783.

Username

Password: K7qzzhq2

## Supporting information

Mouery supplemental figures 1-6

## Acknowledgments and funding sources

We thank Thomas Webb and Natalie Barker-Krantz for technical assistance. We thank Jeffrey Jones for managerial assistance and the Cook lab, Dr. Michael Emanuele, and Dr. Robert Duronio for helpful discussion. The RPE1-hTERT cells containing inducible mTurq-EMI1 and the pCW-puro_mTurq-EMI1-WT plasmid were a gift from Dr. R. Medema. The pTRIPZ-puro_2xFLAG-2xSTREP-Emi1-WT and pTRIPZ-puro_2xFLAG-2xSTREP-Emi1-ΔF-box plasmids were gifts from Dr. M. Pagano.

BL Mouery was supported by NIH/NIGMS 5T32GM007092 and 1T32GM135128 as well as by NIH/NCI 5F31CA268866-02. The project was supported by grants from the NIH: R35GM141833 to JGC. The UNC Flow Cytometry Core Facility (RRID:SCR_019170) is supported in part by P30 CA016086 Cancer Center Core Support Grant to the UNC Lineberger Comprehensive Cancer Center. Research reported in this publication was supported in part by the North Carolina Biotech Center Institutional Support Grant 2017-IDG-1025 and by the National Institutes of Health 1UM2AI30836-01. The content is solely the responsibility of the authors and does not necessarily represent the official views of the National Institutes of Health. This research is based in part upon work conducted using the UNC Proteomics Core Facility, which is supported in part by NCI Center Core Support Grant (2P30CA016086-45) to the UNC Lineberger Comprehensive Cancer Center

## Figure Legends

**Figure S1. Gating strategy used to quantify cell cycle distribution.**

Cells were treated as described in figure 1B. Shown is an example gating strategy to determine the percentage of cells in either G1, S, or G2/M phase.

**Figure S2. Titration of EMI1 expression.**

**(A)** Immunoblot of RPE1 cells treated with 1 µM CDK4/6i for 48 hours with increasing concentrations of doxycycline to induce EMI1 expression during the final 24 hours as indicated.

**(B)** Quantification of EMI1 over-expression from lysates in (A). The ectopic EMI1 (mTurq-EMI1) at each doxycycline concentration was first normalized to total protein (Ponceau S stain) and then normalized to the amount of endogenous EMI1 in the asynchronous/untreated sample (lane 1). Dashed line displays a value of 1 where the ectopic EMI1 over-expression is equivalent to the amount of endogenous EMI1 in the proliferating control sample.

**(C)** Cell cycle distribution of RPE1 cells treated as in (A). One sample was collected after 24 hours of CDK4/6i to show that cells were fully arrested at the time of EMI1 induction.

1 biological replicate was performed.

**Figure S3. APC/C inhibition in quiescent cells.**

**(A)** Cell cycle distribution of RPE1 cells contact inhibited for 120 hours ± EMI1 expression during the final 72 hours as indicated. One sample was collected after 48 hours of contact inhibition to show that cells were fully arrested at the time of EMI1 induction.

**(B)** Cell cycle distribution of RPE1 cells serum starved for 48 hours ± EMI1 expression during the final 24 hours as indicated. One sample was collected after 24 hours of serum starvation to show that cells were fully arrested at the time of EMI1 induction.

For panel A: one representative example of 2 biological replicates is shown. One biological replicate of panel B was performed.

**Figure S4. EMI1 expression bypasses CDK4/6i-induced G1 arrest independently of its F-box domain.**

**(A)** Cell cycle distribution of RPE1 cells treated with 1 µM CDK4/6i for 48 hours ± expression of either WT EMI1 or ΔFbox EMI1 during the final 24 hours as indicated. One sample was collected after 24 hours of CDK4/6i to show that cells were fully arrested at the time of EMI1 induction.

**(B)** Immunoblot of lysates from (A). An untreated proliferating sample from each cell line (lane 1 and 5) was included as a control. Geminin blot shows that ΔFbox EMI1 retains its ability to inhibit the APC/C.

For cell cycle and immunoblots, one representative of 2 biological replicates is shown.

**Figure S5. E2F1 knockdown partially impairs arrest bypass.**

**(A)** Cell cycle distribution of RPE1 cells treated with 1 µM CDK4/6i for 20 hours followed by transfection with either control siRNA (“CTRL”) or siRNA targeting E2F1 (“siE2F1”) for 8 hours prior to 24 hours ± EMI1 expression as indicated (52 hours total; cells remain in CDK4/6i for the entire duration). One sample from each siRNA transfection was collected after 28 hours CDK4/6i to show that cells were fully arrested at the time of EMI1 induction.

**(B)** Immunoblot of lysates from (A). An untreated proliferating sample (lane 1) was included as a control.

For cell cycle and immunoblots, one representative of 2 biological replicates is shown.

**Figure S6. Two independent siRNAs targeting cyclin A or B prevent CDK4/6-inhibited cells from escaping arrest.**

**(A)** Cell cycle distribution of asynchronous RPE1 cells transfected with either control siRNA (“CTRL”) or siRNA targeting cyclin A (“siCCNA2-1”) or cyclin B (“siCCNB1-1”) for 48 hours.

**(B)** Immunoblot from lysates in (A).

**(C)** Cell cycle distribution of asynchronous RPE1 cells transfected with either control siRNA (“CTRL”) or siRNA targeting cyclin A (“siCCNA2-2”) or cyclin B (“siCCNB1-2”) for 48 hours. The siRNAs targeting cyclin A or cyclin B are distinct sequences from those used in (A).

**(D)** Immunoblot from lysates in (C).

**(E)** Cell cycle distribution of RPE1 cells treated with 1 µM CDK4/6i for 24 hours followed by transfection with either control siRNA (“CTRL”) or siRNA targeting either cyclin A (“siCCNA2-2”) or cyclin B (“siCCNB1-2”) for 24 hours prior to 24 hours ± EMI1 expression as indicated (72 hours total; cells remain in CDK4/6i for the entire duration). One sample from each siRNA transfection was collected after 48 hours of CDK4/6i to show that cells were fully arrested at the time of EMI1 induction. The siRNAs targeting cyclin A or cyclin B are distinct sequences from those used in figure 4.

**(F)** Immunoblot of lysates from (E). An untreated proliferating sample (lane 1) was included as a control.

For cell cycle and immunoblots, one representative example is shown. Panels A-B are representative of 2 biological replicates. For panels C-F: 1 biological replicate was performed.

