## Supplementary material for "APC/C prevents non-canonical order of cyclin/CDK activity to maintain CDK4/6 inhibitor-induced arrest": Mouery supplemental figures 1-6

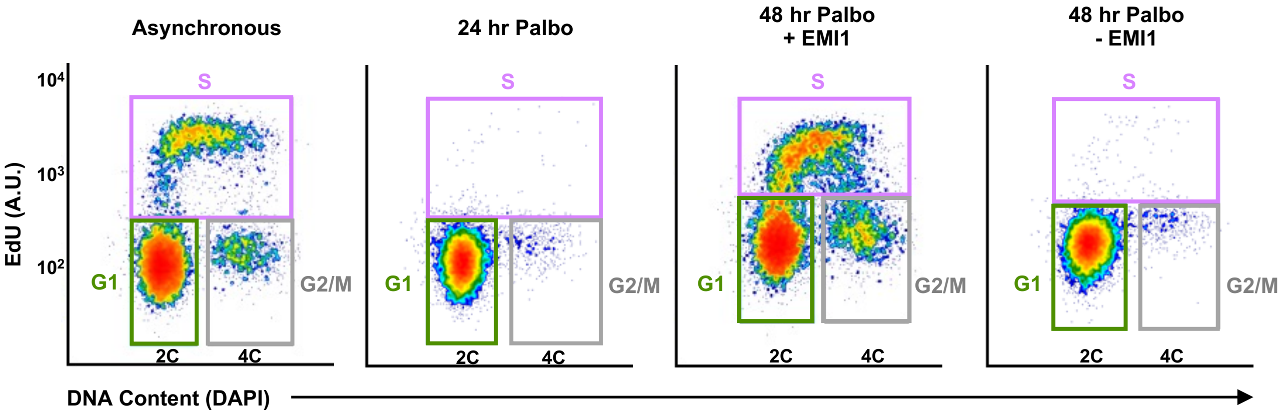


**Figure S1. Gating strategy used to quantify cell cycle distribution.**
Cells were treated as described in figure 1B. Shown is an example gating strategy to determine the percentage of cells in either G1, S, or G2/M phase.

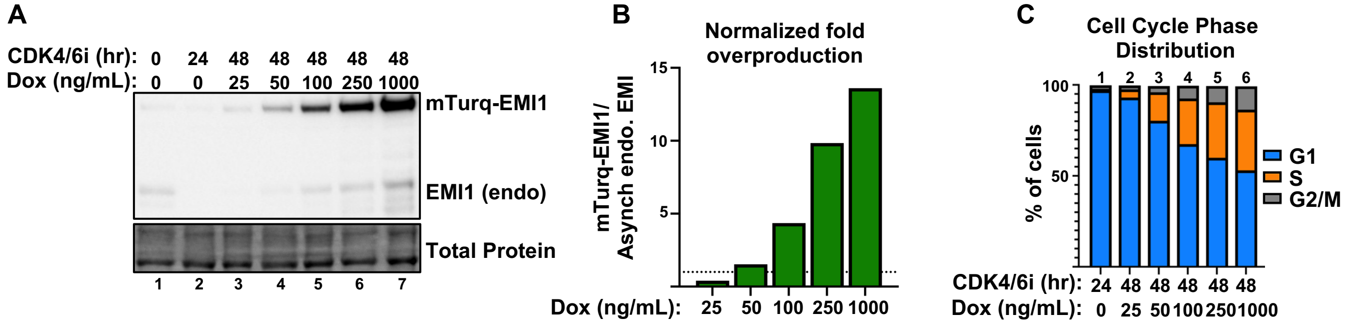


**Figure S2. Titration of EMI1 expression.**

**(A)** Immunoblot of RPE1 cells treated with 1 μM CDK4/6i for 48 hours with increasing concentrations of doxycycline to induce EMI1 expression during the final 24 hours as indicated. **(B)** Quantification of EMI1 over-expression from lysates in (A). The ectopic EMI1 (mTurq-EMI1) at each doxycycline concentration was first normalized to total protein (Ponceau S stain) and then normalized to the amount of endogenous EMI1 in the asynchronous/untreated sample (lane 1). Dashed line displays a value of 1 where the ectopic EMI1 over-expression is equivalent to the amount of endogenous EMI1 in the proliferating control sample. **(C)** Cell cycle distribution of RPE1 cells treated as in (A). One sample was collected after 24 hours of CDK4/6i to show that cells were fully arrested at the time of EMI1 induction.

1 biological replicate was performed.

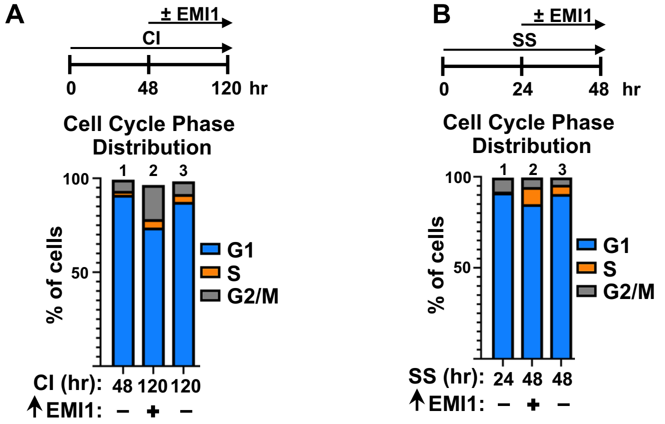


**Figure S3. APC/C inhibition in quiescent cells.**

**(A)** Cell cycle distribution of RPE1 cells contact inhibited for 120 hours ± EMI1 expression during the final 72 hours as indicated. One sample was collected after 48 hours of contact inhibition to show that cells were fully arrested at the time of EMI1 induction. **(B)** Cell cycle distribution of RPE1 cells serum starved for 48 hours ± EMI1 expression during the final 24 hours as indicated. One sample was collected after 24 hours of serum starvation to show that cells were fully arrested at the time of EMI1 induction.

For panel A: one representative example of 2 biological replicates is shown. One biological replicate of panel B was performed.

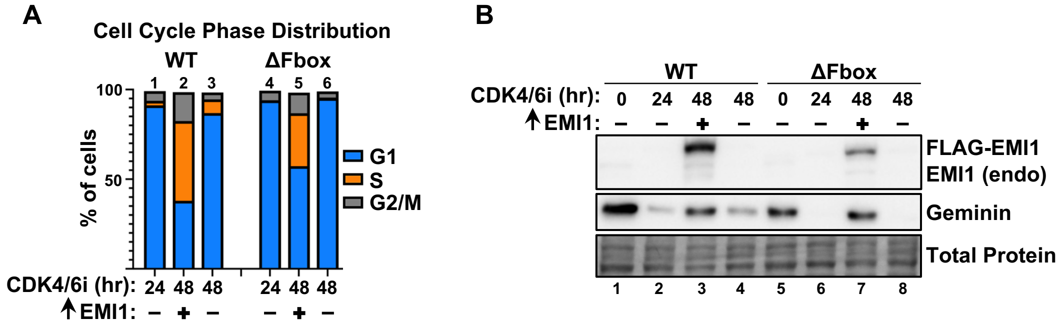


**Figure S4. EMI1 expression bypasses CDK4/6i-induced G1 arrest independently of its F-box domain.**
**(A)** Cell cycle distribution of RPE1 cells treated with 1 μM CDK4/6i for 48 hours ± expression of either WT EMI1 or ΔFbox EMI1 during the final 24 hours as indicated. One sample was collected after 24 hours of CDK4/6i to show that cells were fully arrested at the time of EMI1 induction. **(B)** Immunoblot of lysates from (A). An untreated proliferating sample from each cell line (lane 1 and 5) was included as a control. Geminin blot shows that ΔFbox EMI1 retains its ability to inhibit the APC/C.

For cell cycle and immunoblots, one representative of 2 biological replicates is shown.

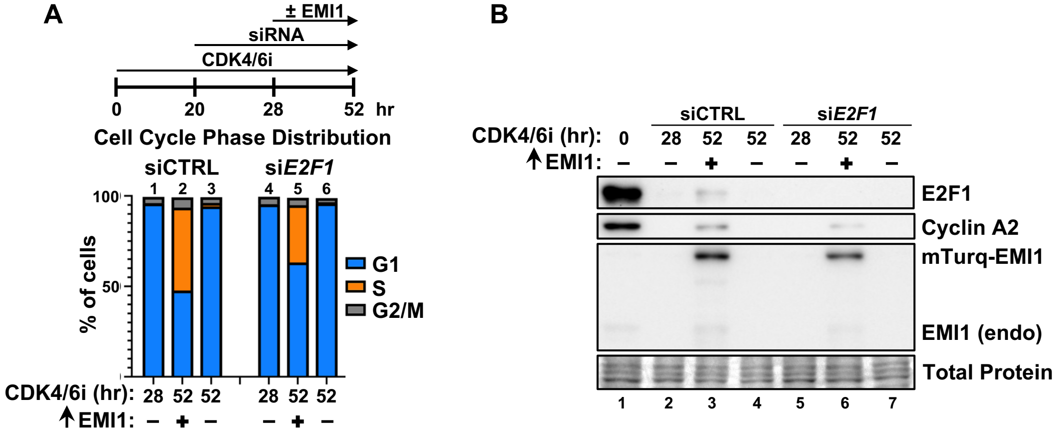


**Figure S5. E2F1 knockdown partially impairs arrest bypass.**
**(A)** Cell cycle distribution of RPE1 cells treated with 1 μM CDK4/6i for 20 hours followed by transfection with either control siRNA ("CTRL") or siRNA targeting E2F1 ("siE2F1") for 8 hours prior to 24 hours ± EMI1 expression as indicated (52 hours total; cells remain in CDK4/6i for the entire duration). One sample from each siRNA transfection was collected after 28 hours CDK4/6i to show that cells were fully arrested at the time of EMI1 induction. **(B)** Immunoblot of lysates from (A). An untreated proliferating sample (lane 1) was included as a control.
For cell cycle and immunoblots, one representative of 2 biological replicates is shown.

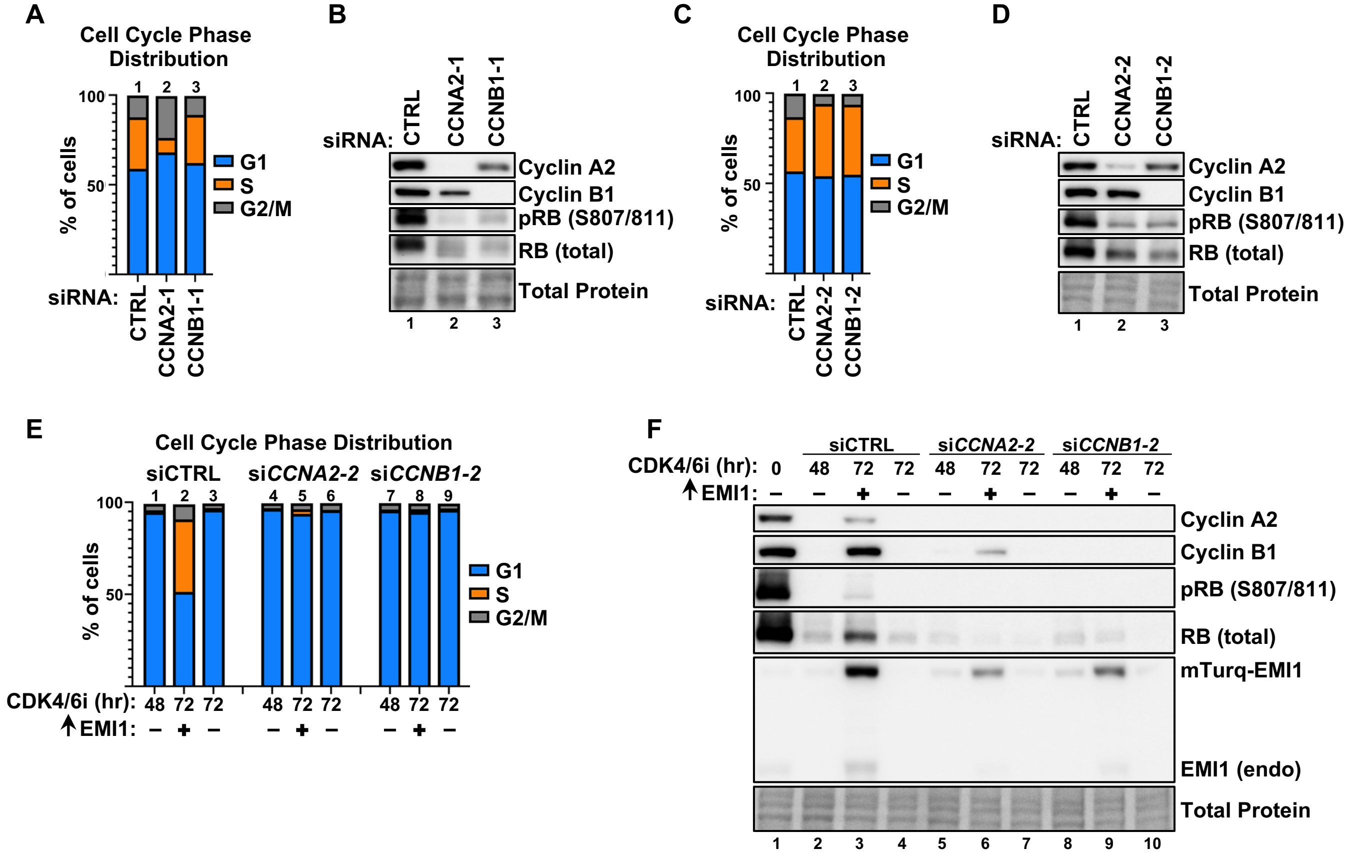


**Figure S6. Two independent siRNAs targeting cyclin A or B prevent CDK4/6-inhibited cells from escaping arrest.**
**(A)** Cell cycle distribution of asynchronous RPE1 cells transfected with either control siRNA ("CTRL") or siRNA targeting cyclin A ("siCCNA2-1") or cyclin B ("siCCNB1-1") for 48 hours. **(B)** Immunoblot from lysates in (A). **(C)** Cell cycle distribution of asynchronous RPE1 cells transfected with either control siRNA ("CTRL") or siRNA targeting cyclin A ("siCCNA2-2") or cyclin B ("siCCNB1-2") for 48 hours. The siRNAs targeting cyclin A or cyclin B are distinct sequences from those used in (A). **(D)** Immunoblot from lysates in (C). **(E)** Cell cycle distribution of RPE1 cells treated with 1 μM CDK4/6i for 24 hours followed by transfection with either control siRNA ("CTRL") or siRNA targeting either cyclin A ("siCCNA2-2") or cyclin B ("siCCNB1-2") for 24 hours prior to 24 hours ± EMI1 expression as indicated (72 hours total; cells remain in CDK4/6i for the entire duration). One sample from each siRNA transfection was collected after 48 hours of CDK4/6i to show that cells were fully arrested at the time of EMI1 induction. The siRNAs targeting cyclin A or cyclin B are distinct sequences from those used in figure 4. **(F)** Immunoblot of lysates from (E). An untreated proliferating sample (lane 1) was included as a control.
For cell cycle and immunoblots, one representative example is shown. Panels A-B are representative of 2 biological replicates. For panels C-F: 1 biological replicate was performed.
